# MAP: A Knowledge-driven Framework for Predicting Single-cell Responses for Unprofiled Drugs

**DOI:** 10.64898/2026.02.25.708091

**Authors:** Jinghao Feng, Ziheng Zhao, Xiaoman Zhang, Mingfei Liu, Jingyi Chen, Xingran Quan, Jian Zhang, Yanfeng Wang, Ya Zhang, Weidi Xie

**Affiliations:** School of Artificial Intelligence, Shanghai Jiao Tong University, Shanghai, China; Shanghai Artificial Intelligence Laboratory, Shanghai, China; Department of Biomedical Informatics, Harvard Medical School, Boston, MA, USA; Department of Pharmaceutical and Artificial-Intelligence Sciences, Shanghai Jiao Tong University School of Medicine, Shanghai, China

## Abstract

Predicting how cells respond to chemical perturbations is one of the goals for building virtual cells, yet experimentally profiled compounds cover only a small fraction of this space. Existing models struggle to generalize to unprofiled compounds, as they typically treat drugs as isolated identifiers without encoding their mechanistic relationships. We present **MAP**, a framework that integrates structured biological knowledge into cellular perturbation modeling and supports zero-shot prediction for small molecules with scarce or absent perturbation profiles. Specifically: (i) we construct **MAP-KG**, a large-scale knowledge graph tailored for cellular perturbation modeling that unifies 14 public resources, spanning 187k drugs, 23k genes, and 694k mechanistic relationships; (ii) we propose a knowledge-driven pre-training strategy that aligns molecular structures, protein sequence features, and textual mechanistic descriptions into a unified embedding space via contrastive learning, producing mechanism-aware and transferable gene and compound embeddings. The resulting knowledge-informed gene and drug representations are then coupled with a pretrained single-cell foundation model to condition perturbation response prediction; (iii) we evaluate MAP under two zero-shot generalization regimes: unseen cell type–drug combinations and the stricter setting of unprofiled drugs, where it improves top-50 DEG Pearson delta correlation by up to +13.3% and +12.2%, respectively, over the strongest baselines across three benchmarks. We further perform pathway-level functional analysis via GSEA for in-silico screening, where MAP predicts coherent, mechanism-consistent programs on unprofiled candidate drugs, and prioritizes 4 of 5 approved anti-cancer drugs in A-549 (non–small cell lung cancer).

## 1 Introduction

Single-cell perturbation profiling is transforming our ability to interrogate causal drug actions by measuring transcriptome-wide responses at cellular resolution [1–5]. By linking chemical interventions to downstream gene programs across heterogeneous cell states, these atlases provide a critical substrate for constructing virtual cells, *i*.*e*., predictive models that forecast cellular responses under novel perturbations and contexts [6–8]. If accurate and generalizable, such models could enable scalable *in silico* screening across large chemical spaces, prioritizing candidate therapeutics and dosing strategies before physical experimentation and thereby reducing the cost and latency of early-stage drug discovery [9].

Despite rapid progress driven by larger perturbation atlases and more expressive models [10–16], robust generalization to *unprofiled* compounds remains a central challenge. In many atlas-scale settings, compounds are treated as categorical identifiers, and perturbation effects are inferred primarily from observed expression profiles [14, 15]. Such representations implicitly place drugs in a latent space where proximity is learned only from co-occurrence in the training atlas, rather than from shared biological mechanisms, such as shared binding modes or similar pathway modulation. As a result, mechanistically related compounds may be encoded as unrelated tokens, limiting extrapolation when a test compound has little or no transcriptional profiling data.

A natural remedy is to condition predictors on drug descriptors such as molecular structure or targets, which can partially support out-of-distribution prediction. However, these descriptors are heterogeneous and incomplete in practice, and they do not, by themselves, provide a unified interface for integrating diverse biological evidence (*e*.*g*., SMILES strings, protein sequences, pathway membership, and free-text mechanism descriptions) into a single mechanism-aware representation. Consequently, current models [10, 12, 17–21] still struggle to systematically transfer such knowledge to the setting that matters most for discovery: predicting transcriptional responses for novel compounds outside the profiled atlas [22].

Here we present **MAP** (Mechanism-Aware Perturbation response predictor), a framework that integrates structured pharmacological knowledge into cellular response prediction. MAP learns mechanism-aware drug and gene representations by aligning molecular structures, protein targets, and mechanistic descriptions in a unified embedding space, and then conditions a perturbation predictor on these knowledge-informed representations. This design enables extrapolation to novel compounds with scarce or absent perturbation profiles, providing a mechanistically grounded route toward *in silico* screening at scale.

Specifically, we make the following contributions.

**On knowledge injection**, we start by constructing MAP-KG (Figure 1a), a perturbation-oriented biomedical knowledge graph that consolidates evidence from 14 public resources. MAP-KG links 187,089 drugs and 22,924 genes through 694,246 mechanistic relations, and leverages text (*e*.*g*., mechanisms of action and functional descriptions) as a semantic bridge connecting heterogeneous attributes such as SMILES strings and protein sequences. Building on MAP-KG, we develop modality-specific encoders and align them via contrastive pre-training to map multi-source evidence into a shared embedding space (Figure 1b).

**Figure 1.**
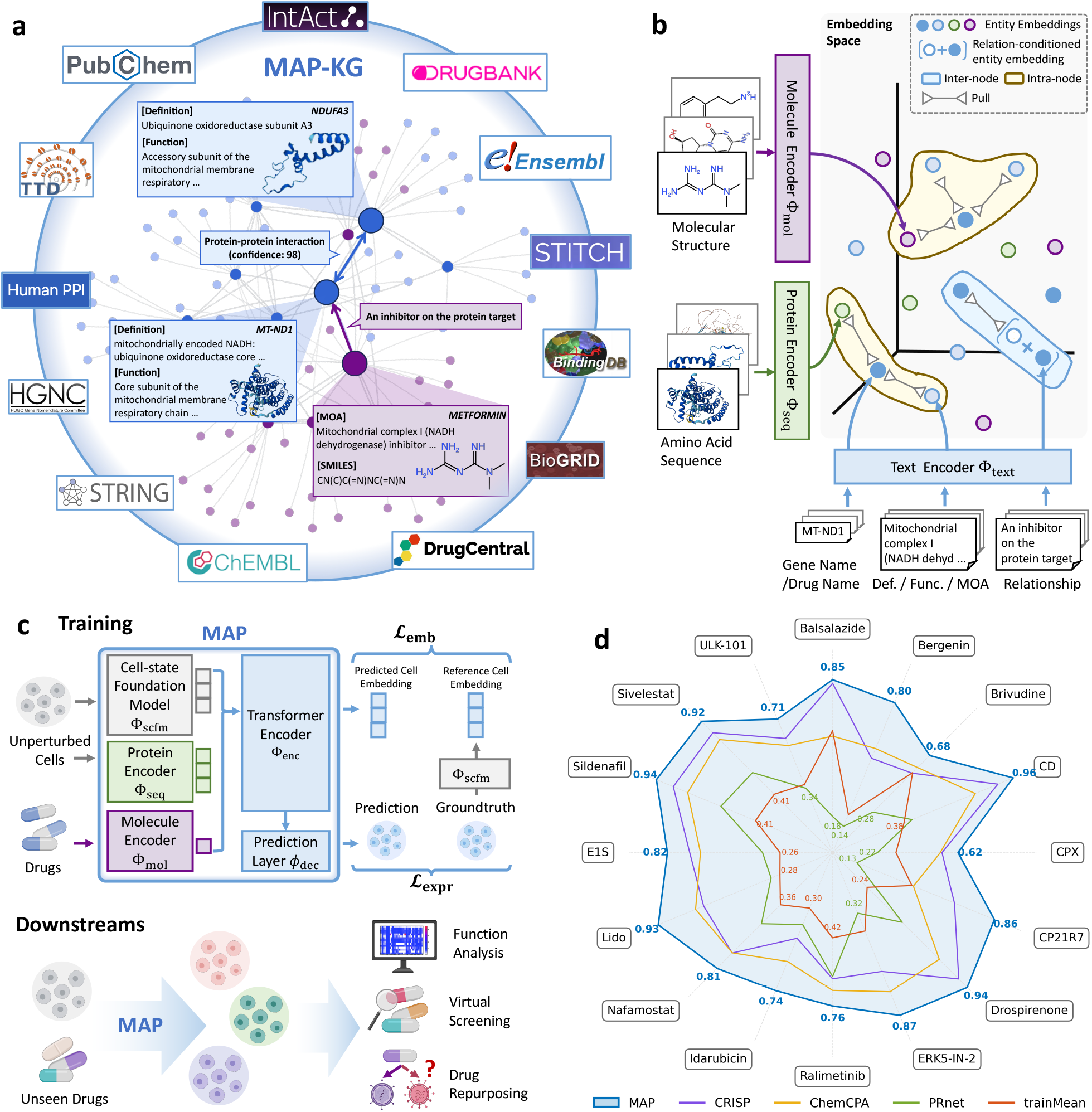
Overview of MAP-KG and MAP. **(a)** We curate 14 authoritative public resources and construct a large-scale biomedical knowledge graph, comprising 187,089 drugs, 22,924 genes and 694,246 triplet relationships. **(b)** We adopt modality-specific encoders for knowledge encoding, and conduct knowledge-driven pre-training to align drug and genes entities with biological mechanism in the representation geometry; **(c)** MAP integrates the knowledge-informed drug and gene representation with a single-cell foundation model to guide perturbation response prediction. Consequently, MAP can generalize to unprofiled drugs without any transcriptional perturbation data, facilitating applications like functional analysis, virtual screening, and drug repurposing; **(d)** MAP demonstrates state-of-the-art predictive performance across a variety of unprofiled drugs. E1S: Estrone sulfate; CPX: Ciclopirox; CD: Carbidopa; Lido: Lidocaine; Ralimetinib: Ralimetinib dimesylate. **(c)** is created with BioRender.

**On model development**, we propose a knowledge-driven pre-training strategy that learns transferable drug and gene representations from MAP-KG by jointly aligning multi-modal attributes and enforcing relation-level consistency. These representations provide a mechanistic prior that moves beyond treating compounds as independent identifiers. We integrate the resulting knowledge encoders into a perturbation predictor built on a single-cell foundation model, combining control-state cell embeddings with knowledge-informed drug and gene embeddings to forecast perturbed transcriptomes (Figure 1c).

**On experiment evaluation**, we validate MAP along multiple axes: (i) for zero-shot compositional generalization to unseen cell type–drug combinations, MAP improves top-50 DE Pearson delta correlation by 13.3% on Tahoe-100M and 8.7% on SciPlex3, and improves direction accuracy by 6.7% on the OP3 benchmark; (ii) for zero-shot prediction of unprofiled drugs (no perturbation profiles observed during training), MAP improves top-50 DE Pearson delta correlation by 12.2% (Tahoe-100M), 8.2% (OP3), and 10.4% (SciPlex3); (iii) using pathway-level GSEA readouts, MAP recovers coherent, mechanism-consistent programs and supports *in silico* screening in A549 (NSCLC), prioritizing 4 of 5 approved anticancer drugs within the top 15 among 58 held-out compounds; (iv) finally, through progressive experiments, we show that scaling mechanistic knowledge provides a complementary axis to data and model capacity, yielding additional gains in zero-shot generalization to unprofiled compounds with substantially lower resource costs.

## 2 Results

We present MAP, a mechanism-aware predictor of transcriptional responses to chemical perturbations that leverages structured biomedical evidence to generalize to compounds with limited or no profiling data. We evaluate MAP in two zero-shot regimes: (i) unseen cell line–drug combinations and (ii) entirely unprofiled drugs, and assess performance using both gene-level metrics and pathway-level GSEA readouts to test functional coherence and utility for *in silico* screening.

### 2.1 Benchmark Description

#### Evaluation settings

We evaluate MAP under two zero-shot protocols of increasing stringency:

##### (i) Zero-shot compositional generalization (unseen cell line–drug combinations)

This setting probes context transfer, where a drug may be observed in the atlas but not in a particular cellular context. We hold out a subset of cell line–drug pairs during training and evaluate performance on these unseen combinations.

##### (ii) Zero-shot prediction for unprofiled drugs (unseen compounds)

This setting mimics virtual screening for candidates with no perturbational measurements. We remove *all* perturbation profiles of a subset of drugs from training and evaluate on these held-out compounds. To prevent information leakage through auxiliary knowledge, we also exclude the held-out drugs from MAP-KG during the knowledge-driven pre-training, so that inference for an unprofiled compound relies only on its available attributes (*e*.*g*., molecular structure and associated annotations) rather than any atlas-derived transcriptional evidence.

Beyond gene-level metrics, we test whether zero-shot predictions preserve coherent, mechanism-relevant transcriptional programs. Under the unprofiled-drug split, we apply GSEA to predicted and observed responses and evaluate pathway-level agreement. We additionally assess *in silico* screening utility on 58 candidate drugs in A549 (non–small cell lung cancer).

##### Datasets

We benchmark MAP across four large-scale datasets covering distinct biological contexts: (i) **Tahoe-100M** [23], a giga-scale atlas spanning 50 cancer cell lines and 379 drugs. To provide a proof-of-concept evaluation while keeping computation tractable, we restrict training and validation to six cell lines, spanning diverse tissues and cancer types. (ii) **OP3** [24], covering 144 compounds across six immune cell types; and (iii) **SciPlex3** [2], profiling 187 compounds across three cancer cell lines. Together, these datasets span diverse cellular contexts and perturbation libraries, enabling systematic evaluation of drug generalization from large cancer atlases to immune benchmarks.

##### Metrics

We evaluate response prediction at the pseudobulk level using three complementary metrics: **(i) Pearson delta correlation**, defined as the Pearson correlation between predicted and observed perturbation-induced expression changes (log fold-changes relative to control); **(ii) Direction accuracy**, the fraction of genes whose predicted sign of regulation (up/down relative to control) matches the observed sign; and **(iii) Perturbation discrimination score**, which quantifies whether predicted responses remain separable across different perturbations. Formal definitions are provided in Section A.3. To mitigate single-cell measurement noise, all evaluations are performed at pseudobulk resolution by aggregating a fixed number of cells within each experimental context; following [15], each pseudobulk profile is constructed by randomly sampling a fixed number of cells from the same condition. We compute the above three metrics on two canonical gene sets: top-2000 highly variable genes (HVG) and top-50 differentially expressed genes (DEG). Note that perturbation discrimination score is calculated only on HVG to ensure gene set consistency across perturbations. For each experiment, we perform five independent runs and report 95% confidence intervals.

##### Baseline models

We compare MAP against state-of-the-art perturbation prediction baselines, including models designed for compositional generalization across contexts and models that condition on molecular structure. Specifically, we include CRISP [19], a scFM-assisted model for zero-shot prediction on unseen cell type–perturbation combinations; chemCPA [10], a compositional perturbation autoencoder that injects drug representations into a latent basal cell state; PRnet [18], a deep generative model that conditions on molecular structure and control-state expression to predict responses to unseen compounds. We also include a competitive linear baseline, trainMean [22, 25], which predicts perturbation effects using mean responses estimated from the training set.

### 2.2 Zero-shot Generalization Across Unseen Cell Line–Drug Combinations

We first assess zero-shot compositional generalization to unseen cell line–drug combinations. In this setting (Figure 2a), we split at the *pair* level: for each cell line, we hold out 5% of drugs *only in that cell line* for testing, while ensuring that the held-out drug sets are non-overlapping across cell lines. Consequently, every test drug remains observed in the training set in other cellular contexts, but its response in the held-out cell line must be inferred without paired supervision. This protocol reflects a repurposing-like scenario in which a compound profiled in some contexts is evaluated in a new cellular setting.

**Figure 2.**
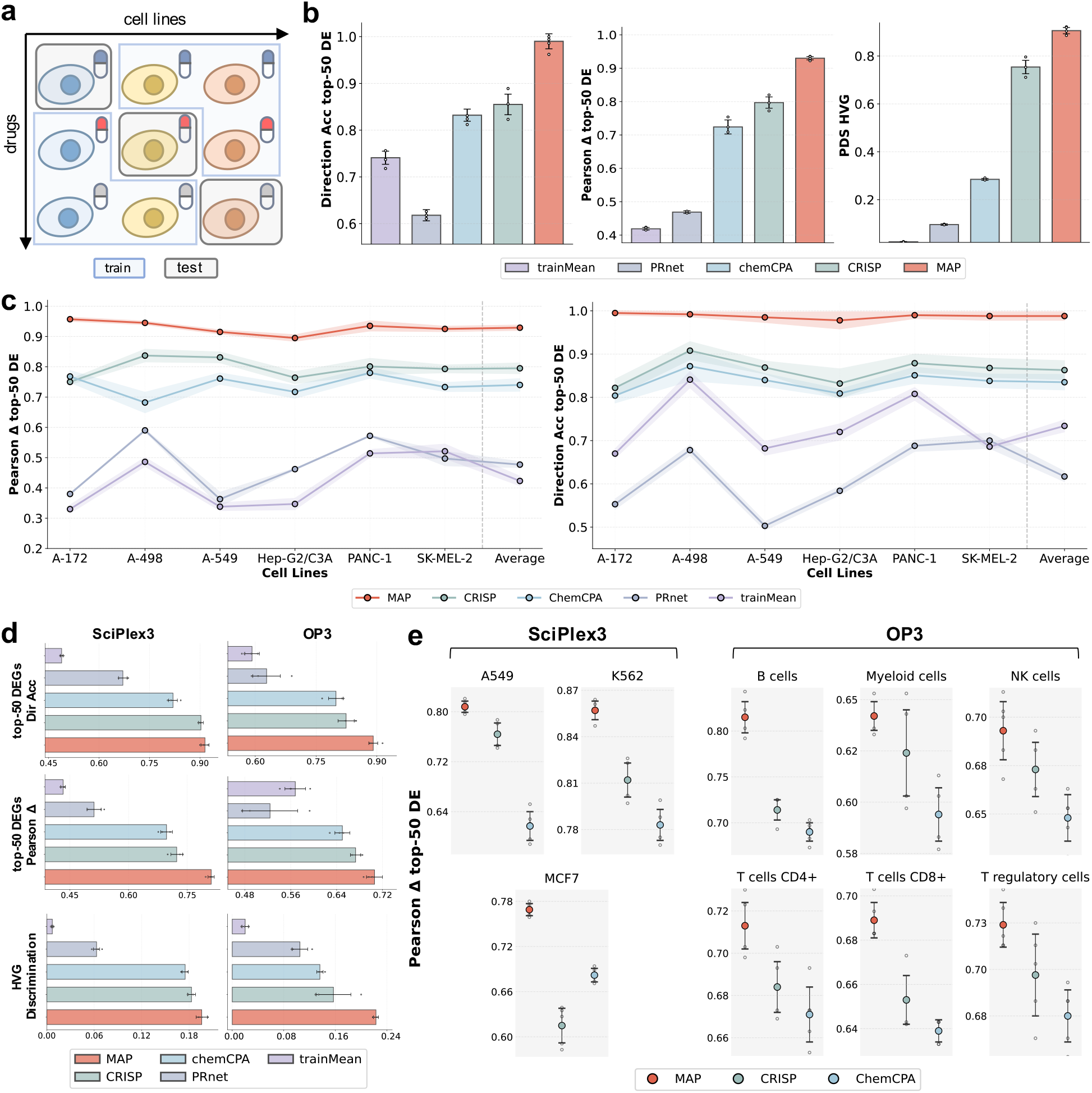
Zero-shot generalization performance on unseen cell line-drug combinations. **(a)** Illustration of zero-shot generalization to unseen cell type-drug pairings. For each cell type, we hold out 5% of drugs as unseen, and the heldout drug sets are non-overlapping across cell types. **(b)** Performance comparison between MAP and baselines on 6 cell lines selected from Tahoe-100M dataset. Three key evaluation metrics are shown. The whiskers represent 95% confidence interval (CI). Five independent data points are shown. **(c)** Cell typespecific performance on Tahoe100M. We report top50 DEG Pearson delta correlation and direction accuracy; shaded bands indicate 95% confidence intervals. **(d)** Performance comparison between MAPand baselines on SciPlex3 and OP3 datasets. Three key evaluation metrics are shown. **(e)** Cell-typespecific performance on SciPlex3 and OP3 datasets. For each cell type, we present the results of the top3 performing methods.

#### Overall performance on Tahoe-100M

As shown in Figure 2b, on Tahoe-100M, MAP significantly outperforms all baselines across metrics, achieving relative improvements over the best-performing baseline (CRISP) of 13.5% in top-50 DEG direction accuracy, 13.3% in top-50 DEG Pearson delta correlation, and 15.2% in the HVG perturbation discrimination score (Details are provided in Supplementary Table S1). These gains indicate that MAP more faithfully matches both the magnitude of perturbation effects (higher Pearson delta correlation) and the direction of regulation (higher direction accuracy), while producing more separable perturbation signatures in the predicted space (higher HVG discrimination score). Together, the results suggest that mechanism-informed compound representations improve transfer of perturbational effects across cellular contexts.

#### Cell line-specific performance on Tahoe-100M

We next stratify performance by cell line (Figure 2c). Across six diverse cancer cell lines spanning multiple tissues and disease contexts, MAP achieves consistently strong results: top-50 DEG Pearson delta correlation reaches ≥ 0.95 on A-172 and remains close to 0.90 on the lowest-performing Hep-G2/C3A line. Direction accuracy is similarly high and stable across cell lines, indicating robust recovery of gene-wise up- and down-regulation. CRISP and chemCPA also improve substantially over the linear trainMean baseline, but their gains are smaller and exhibit greater variability across cell lines than MAP(Details are provided in Supplementary Table S2).

#### Generalization across datasets

To test robustness beyond Tahoe-100M, we repeat the same unseen-combination protocol on **SciPlex3** and the **OP3** benchmark, which contribute 28 and 42 non-overlapping cell type–drug combinations, respectively. For these datasets, we use the same training and evaluation protocol as in Tahoe-100M. As shown in Figure 2d, MAP consistently improves top-50 DEG direction accuracy, top-50 DEG Pearson delta correlation, and HVG perturbation discrimination score on both datasets, indicating reliable zero-shot inference across experimental settings and extending from cancer cell lines to primary immune cell types. We note that HVG perturbation discrimination score values are relatively low for all methods on SciPlex3 and OP3, which may reflect weaker perturbation separability arising from dataset-specific protocol differences and experimental configurations.

#### Cell type-specific analysis on SciPlex3 and OP3

Finally, we examine cell type-specific performance (Figure 2e). On SciPlex3, MAP performs consistently strong across the three cell lines, achieving a top-50 DEG Pearson delta correlation of 0.857 on K562. On OP3, MAP similarly maintains strong performance across all six immune cell types; even in myeloid cells, where correlations are lower for all models, MAP reaches 0.642, indicating meaningful agreement between predicted and observed perturbation effects (Details are provided in Supplementary Table S3).

### 2.3 Zero-shot Generalization to Unprofiled Drugs

We next evaluate MAP in a more challenging regime: zero-shot prediction for *unprofiled drugs*, where test compounds are entirely absent from perturbation training data (Figure 3a). This setting probes whether the model can infer perturbation effects from molecular attributes (*e*.*g*., chemical structure), without access to any transcriptional response measurements for the test compounds in any cell type. We hold out 5% of drugs as the test set and remove all corresponding perturbation profiles from training. To avoid information leakage, we additionally exclude the held-out drug entities (and their aliases, if applicable) from MAP-KG during knowledge pre-training.

**Figure 3.**
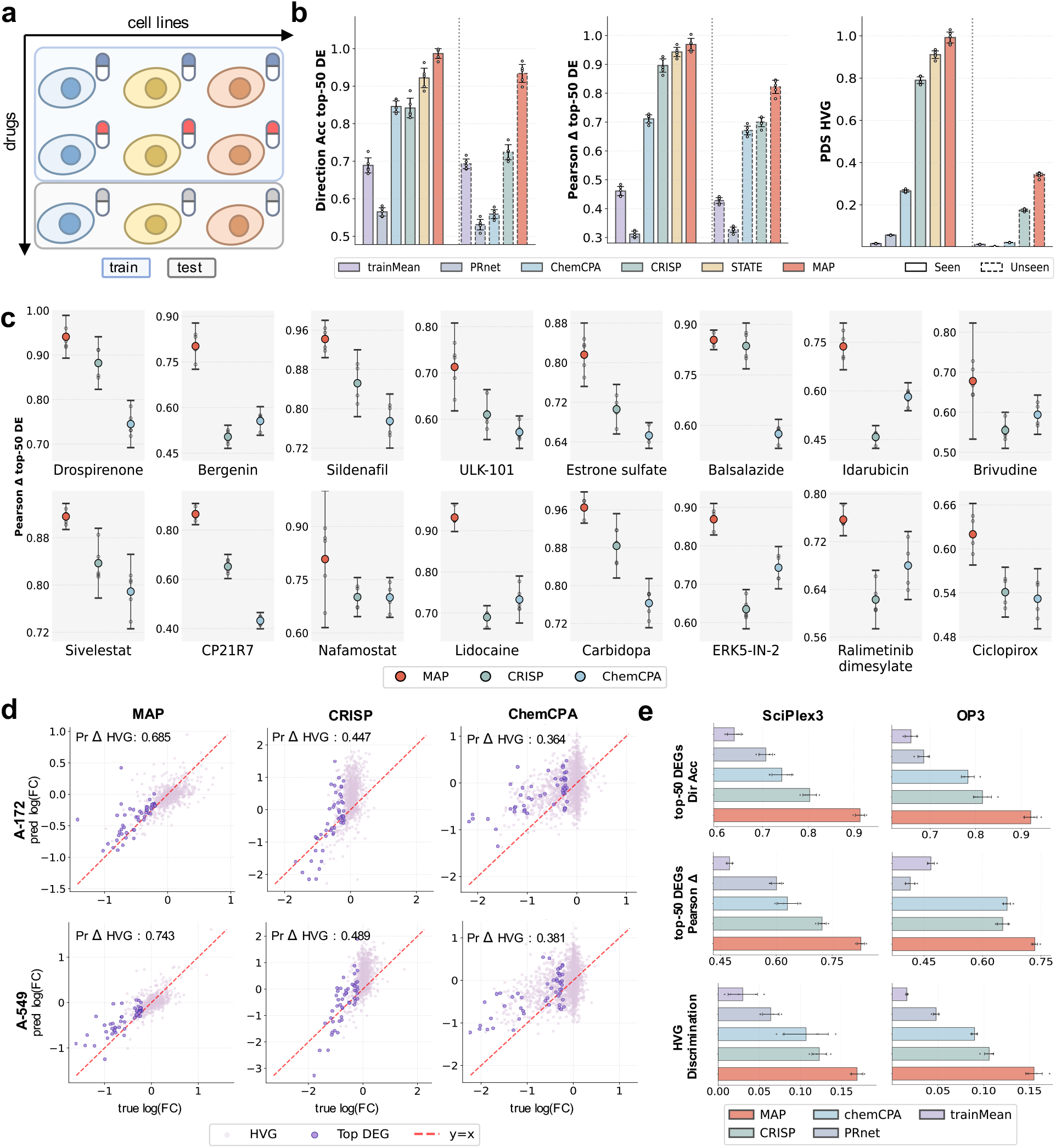
Unprofiled drug generalization performance of MAP. **(a)** Illustration of drug generalization scenario, with 5% drugs withheld during training as unseen testing drugs. **(b)** Average performance comparison between MAP and baseline models on 6 cell lines selected from Tahoe-100M dataset. Three evaluation metrics are shown. The whiskers represent 95% confidence interval (CI), while five independent data points are shown by small dots. **(c)** Compound-specific analysis of top 50 DE genes’ Pearson delta correlation, where 95% CI and five data points are shown. **(d)** Visualization of the single-gene-level perturbation predictions for Idarubicin (hydrochloride) on A-172 and A-549 cell lines. The top 50 DE genes and top 2000 HVG are marked with distinct colors. Pearson delta correlations of HVG are annotated in each panel. **(e)** Performance comparison between MAP and baseline models on OP3 dataset and SciPlex3 dataset. Three evaluate metrics are demonstrated.

#### Overall performance on Tahoe-100M

As shown in Figure 3b, MAP substantially improves over baseline methods across all metrics in the unprofiled-drug regime, with +21.0% top-50 DEG direction accuracy, +12.2% top-50 DEG Pearson delta correlation, and +16.9% HVG perturbation discrimination score. Notably, MAP also improves performance on drugs seen during training, suggesting that knowledge injection not only supports extrapolation to unseen compounds but also strengthens in-distribution modeling by providing a more informative mechanistic prior (Details are provided in Supplementary Tables S4 and S5). For completeness, we also compare against STATE [15], a strong baseline pre-trained on Tahoe-100M.

#### Per-compound performance

We further analyze results at the level of individual held-out compounds (Figure 3c). MAP achieves the highest top-50 DEG Pearson delta correlation on all of 16 test compounds. Across compounds, performance ranges from 0.942 on Sildenafil to 0.620 on Ciclopirox, indicating robust zero-shot prediction despite diverse chemical structures and heterogeneous mechanisms (Details are provided in Supplementary Table S6).

#### Gene-level agreement for representative drugs

To examine gene-level behavior, we visualize single-gene predictions for Idarubicin (hydrochloride) in A-172 and A-549, comparing MAP with CRISP and chemCPA (Figure 3d). MAP shows tighter alignment with the diagonal, indicating closer agreement between predicted and observed gene-wise log fold-changes. Consistently, Pearson delta correlation on HVG is higher for MAP: on A-172, 0.685 versus 0.447 (CRISP) and 0.364 (chemCPA); on A-549, 0.743 versus 0.489 and 0.381, respectively. While a small number of outliers remain, MAP better captures perturbation magnitudes with reduced dispersion and fewer systematic biases than the baselines. These results demonstrate that MAP more faithfully recovers fine-grained transcriptional perturbation patterns at the single-gene level, highlighting superior ability to model expression change across different cellular contexts.

#### Generalization across datasets

Finally, we evaluate unprofiled-drug generalization on SciPlex3 and the OP3 benchmark (Figure 3e). MAP maintains strong gene-level accuracy, achieving an average top-50 DEG Pearson delta correlation of 0.823 across SciPlex3 cell lines and 0.734 across OP3 immune cell types, with consistent gains in direction accuracy and HVG perturbation discrimination score (Details are provided in Supplementary Tables S8 and S9). Together, these results support the robustness of MAP for zero-shot prediction on unprofiled compounds across datasets and heterogeneous cellular contexts.

### 2.4 Functionally Interpretable Pathway Responses

To improve functional interpretability, we evaluate pathway-level responses by applying gene set enrichment analysis (GSEA) [26] to model-predicted perturbation profiles. This shifts assessment from individual genes to coordinated biological programs that are more directly actionable in biomedical settings. We focus on the A-549 cell line, a widely adopted model of lung adenocarcinoma and non–small cell lung cancer (NSCLC) drug response. We curate an expanded set of 58 candidate drugs from Tahoe-100M, including five drugs that have been approved for use in A-549–related contexts: Afatinib, Osimertinib, Crizotinib, Adagrasib, and Pemetrexed. Following the unprofiled-drug protocol (Section 2.3), we train MAP and perform zero-shot prediction on the held-out candidate compounds.

#### Simulated *in-silico* drug screening

We first conduct a simulated screening experiment on A-549 cell line. We curate a set of disease-relevant pathways whose activity is desirable to suppress, including major oncogenic drivers and core signaling axes implicated in lung adenocarcinoma progression. Following [19], for each drug we compute pathway enrichment scores from the predicted transcriptional response and aggregate them across the curated pathways to obtain a drug-level *downregulation score*, where lower scores indicate stronger predicted suppression and therefore more promising candidates. Figure 4a reports the drug ranking by the inverse downregulation score. Four clinically approved lung cancer drugs [27–32] appear among the top 14 candidates; notably, Adagrasib and Afatinib rank second and fourth among all held-out compounds, respectively, despite having no perturbation profiles available during training. We further quantify retrieval performance by computing top-*k* recall of the approved drugs and compare against CRISP and chemCPA. As shown in Figure 4b, MAP achieves consistently higher recall across *k*, suggesting improved utility for virtual screening.

**Figure 4.**
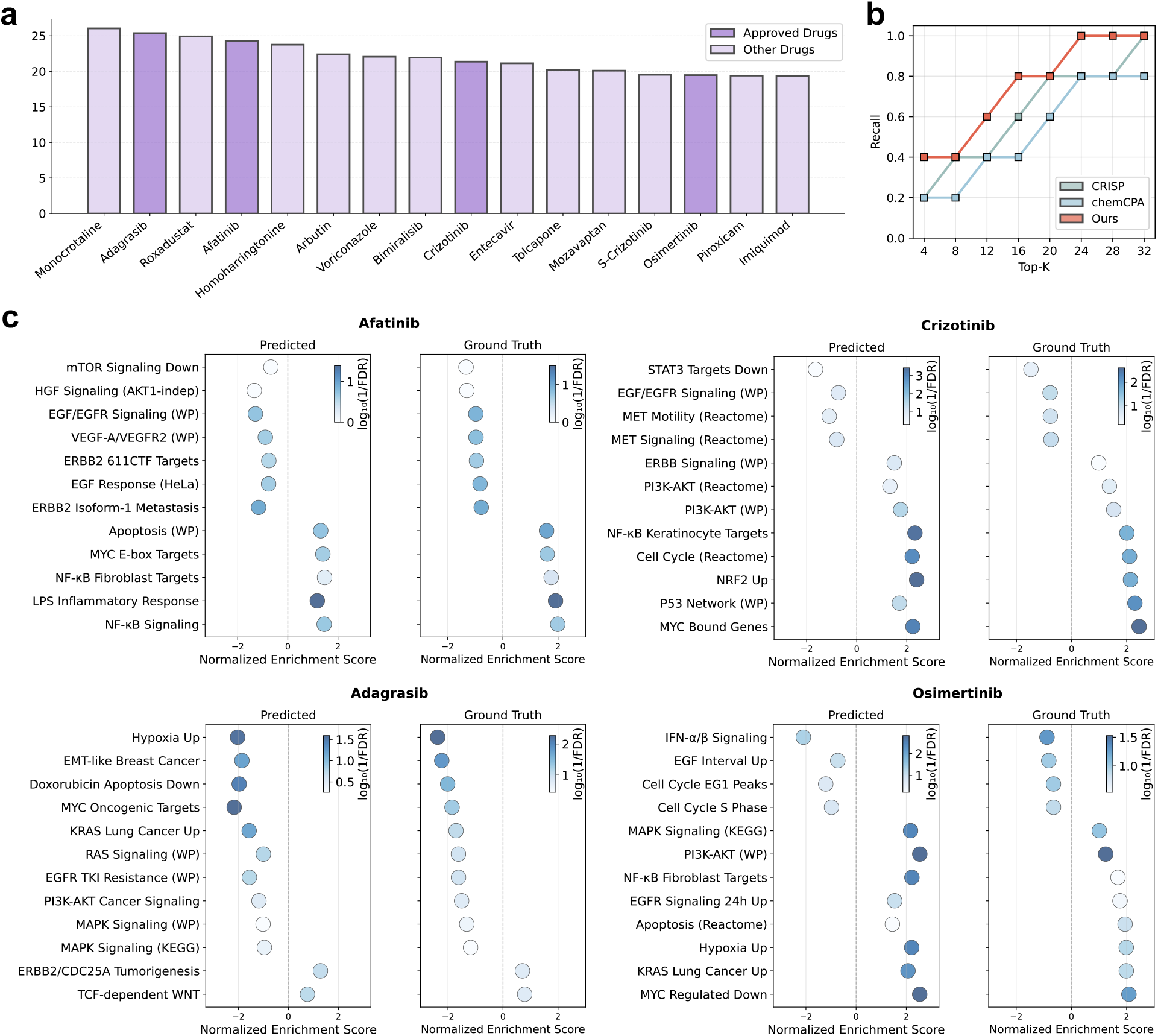
Functionally interpretable pathway responses. **(a)** Virtual drug screening based on aggregated GSEA enrichment scores over significant A-549–relevant pathways, demonstrating top-16 ranked drugs out of 58 unseen drugs. Four clinically approved drugs are prioritized among unseen compounds. **(b)** Top-k recall comparison of clinically approved drugs between MAP, CRISP, and chemCPA, where MAP achieves higher recall at smaller k, indicating more efficient screening. **(c)** Comparison of predicted and experimental GSEA normalized enrichment scores (NES) for key pathways across representative approved drugs. For each drug, twelve gene sets closely related to the A-549 disease-driving context and the drug’s mechanism of action are shown, and MAP closely matches the direction and relative magnitude of pathway regulation observed experimentally.

#### Gene set-level mechanism agreement

To assess whether MAP captures biologically meaningful programs beyond ranking, we perform a fine-grained analysis of pathway regulation. Figure 4c summarizes GSEA results for the four approved drugs that rank highest in the screening experiment. Across these four drugs, the predicted normalized enrichment scores (NES) closely match the ground-truth NES for the highlighted gene sets, correctly recovering both the direction and relative magnitude of pathway regulation. These results indicate that MAP can produce functionally interpretable perturbation predictions at the pathway level that are consistent with established biological mechanisms. For each drug, we additionally examine representative pathways linked to its known mechanism of action and to disease-relevant signaling in A-549 cells; details are provided in the Supplementary Materials Section A.2 and Table S10.

### 2.5 Biological Knowledge Enhances Zero-shot Generalization to Unprofiled Drugs

We posit that structured mechanistic knowledge serves as a critical inductive bias, enabling the model to extrapolate perturbation effects to compounds with limited or absent transcriptional supervision. We validate and dissect the contributions of biological knowledge in unprofiled-drug generalization (Section 2.3), along two complementary axes: (i) On knowledge **scale**, we perform progressive knowledge injection by sequentially incorporating knowledge sources (Figure 5a); (ii) On knowledge **diversity**, we conduct category-wise ablation, selectively withholding specific knowledge categories (Figure 5b). In addition, we visualize drug and gene entities in the latent space (Figure 5c) to demonstrate that knowledge enhancement yields mechanism- and annotation-consistent representation geometry.

**Figure 5.**
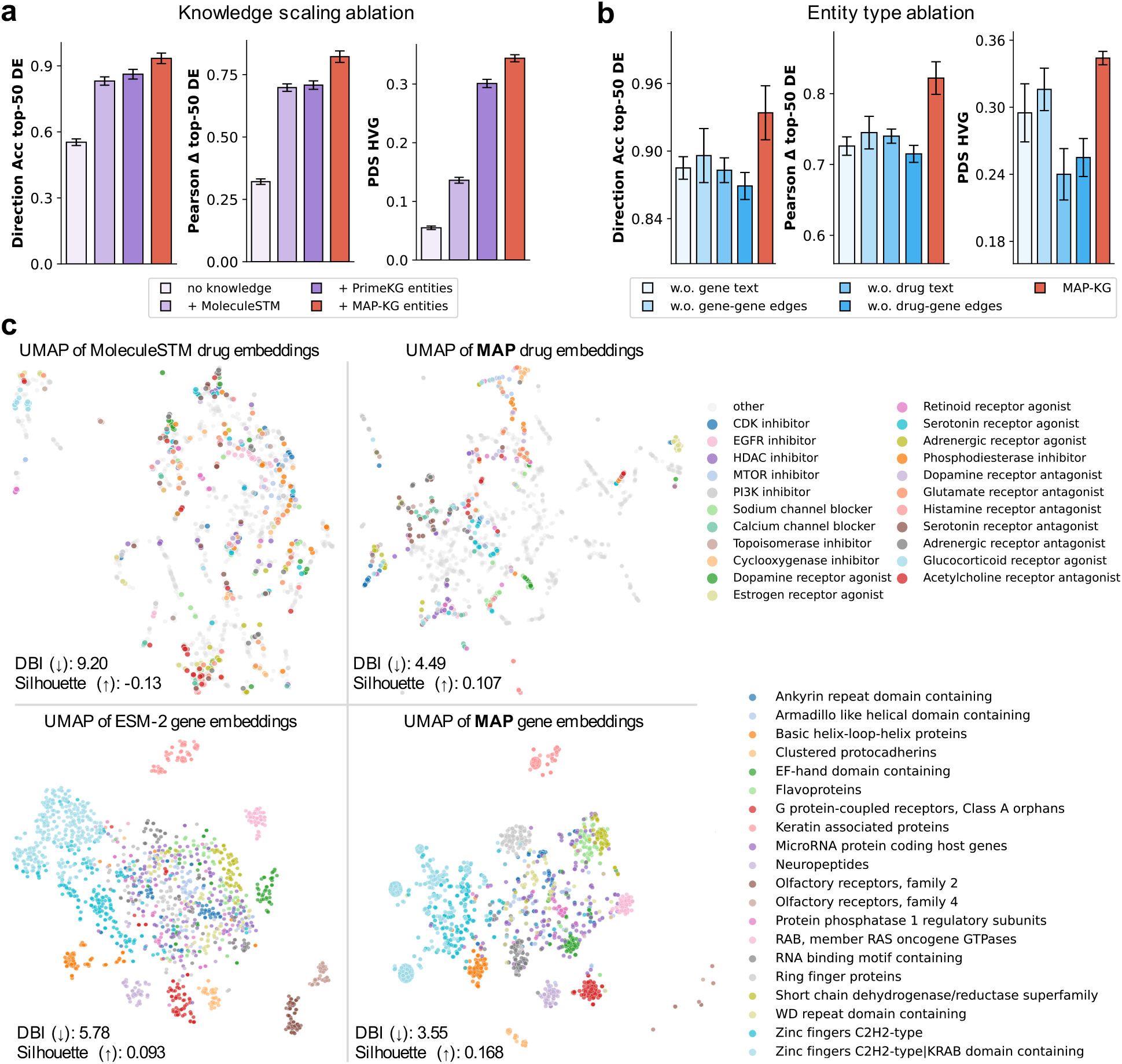
Ablations of knowledge scaling, and visualization of knowledge representations. **(a)** Performance on the unprofiled drug regime as we progressively increase the injected knowledge sources. Knowledge variants are defined as: no knowledge, randomly initialized knowledge encoders; +MoleculeSTM, initialization from a pretrained MoleculeSTM checkpoint; +PrimeKG, further pre-training with PrimeKG entities; +MAP-KG, additional pre-training with MAP-KG entities. Three evaluation metrics and 95% confidence intervals across five runs are reported. **(b)** Ablation of knowledge types by removing one type of entities from MAP-KG at a time. We report the same three metrics under the unprofiled-drug protocol, highlighting the contribution of each knowledge type. **(c)** UMAP visualization of drug embeddings (top) and gene embeddings (bottom) before and after knowledge pre-training. Points are colored by MOA categories for drugs and by gene families for genes, showing clearer mechanism- and function-consistent organization after training. Davies–Bouldin Index(DBI) and Silhouette score are reported.

#### Generalization performance scales with the biological priors

We progressively increase the knowledge data in pre-training, divided into four stages: (i) **No knowledge**, we bypass the knowledge-driven pre-training and use randomly initialized drug embeddings; (ii) **MoleculeSTM**, we skip pre-training and derive drug embeddings from MoleculeSTM [33], an off-the-shelf molecule encoder that provides physicochemical knowledge of chemical structures; (iii) **+PrimeKG**, we adapt the MoleculeSTM with the proposed knowledge-driven pre-training on PrimeKG [34], a knowledge graph centered on drug-disease interactions, containing mechanism-aware knowledge on 7,957 drugs; and (iv) **+MAP-KG**, we continue the pre-training on MAP-KG, a perturbation-oriented knowledge graph of unprecedented scale. As shown in Figure 5a, all three metrics improve monotonically as more knowledge data is included. In particular, adding MoleculeSTM improves top-50 DEG direction accuracy and Pearson Δ correlation by 27.8% and 37.7%, respectively, indicating that drug-side knowledge alone yields rapid gains. Using the full MAP-KG for pre-training further improves over PrimeKG-only pre-training, with additional gains of 7.2%, 11.4%, and 4.3% on top-50 DEG direction accuracy, Pearson delta correlation, and HVG perturbation discrimination score, respectively, underscoring the benefit of scaling knowledge data for generalization performance (Details are provided in Supplementary Table S11).

#### Synergistic knowledge promotes generalization

To investigate the contribution of different biological knowledge, we ablate one knowledge category from MAP-KG at a time, including drug-side textual attributes (*e*.*g*., MOA and therapeutic functions), gene-side attributes (*e*.*g*., amino-acid sequence, textual description of functional annotations), and relational edges (drug–gene mechanistic interactions and gene–gene functional and physical associations). As illustrated in Figure 5b, removing any single category reduces the generalization performance, indicating that the gains are supported by multiple, complementary biological knowledge. Notably, removing drug–gene edges causes the largest drop in both top-50 DEG direction accuracy and Pearson Δ correlation, highlighting the central role of drug–gene mechanistic interactions in cellular perturbation modeling (Details are provided in Supplementary Table S12).

#### Knowledge pre-training yields mechanism-aligned representations

We visualize drug and gene embeddings with UMAP to analyze how multimodal knowledge pre-training reshapes the geometry of entity representations (Figure 5c). We also quantify clustering quality using Davies–Bouldin Index (DBI; lower indicates tighter, better-separated clusters) and Silhouette score (higher indicates better intra-cluster cohesion and inter-cluster separation). For drugs, the pre-trained embeddings show clearer grouping by MOA than the raw MoleculeSTM features, with mechanistically similar compounds forming compact clusters (lower DBI and higher Silhouette), indicating capturing transferable priors among drugs of related mechanism of action. Similarly, for genes, pretrained embeddings exhibit sharper separation by gene family and domain annotations than the ESM-2 embeddings, with lower DBI and higher Silhouette, producing more coherent gene clusters. Together, these patterns suggest that knowledge pre-training promotes mechanism-level information exchange between drug and gene entities, providing a strong basis for transferable, zero-shot perturbation prediction.

## 3 Discussion

This work establishes a novel paradigm that seamlessly bridges mechanism-aware biological knowledge with high-dimensional single-cell readouts, paving the way for more generalizable cellular perturbation modeling. Specifically, we contribute MAP-KG, a perturbation-oriented knowledge graph of unprecedented scale that harmonizes heterogeneous biological knowledge, and MAP, a holistic methodology that transforms these priors into unified drug and gene embeddings and synergizes with existing cell-state foundation models. In the following, we dissect the implications of our work from three perspectives.

### Advancing robust and generalizable artificial-intelligence virtual cells (AIVC)

Building robust AIVCs hinges on the ability to accurately model cellular responses to perturbations. However, a fundamental asymmetry persists: the vastness of the chemical intervention space far outstrips the capacity of experimentally profiling. Despite recent progress, generalization to unprofiled or sparsely observed compounds remains a key bottleneck for existing data-driven paradigms. By introducing comprehensive biological priors, MAP achieves significant advantages in zero-shot perturbation response prediction under unseen cell line–drug combinations (Section 2.2) and unprofiled drugs (Section 2.3). Crucially, the performance superiority is consistently observed across datasets and diverse cellular contexts, confirming that biological knowledge successfully serves as transferable priors even when transcriptional supervision is sparse or absent. By effectively bridging this generalization gap, the paradigm established in this work lays a solid foundation for AIVCs.

### Accelerating therapeutic discovery via biologically faithful virtual screening

The high attrition rate and exorbitant costs of drug discovery necessitate effective *in silico* prioritization strategies. However, virtual screening depends not merely on numerical accuracy at the gene level, but on the faithful prediction of biological functions and regulatory programs. Our simulated screening on the A-549 lung cancer model highlights the translational potential of the proposed approach (Section 2.4). In a strict zero-shot setting, MAP successfully prioritized clinically approved therapies, *e*.*g*., Adagrasib and Afatinib, from a pool of unprofiled candidates, driven by the precise prediction of disease-relevant pathway suppression. By delivering functional interpretable predictions, our framework holds great promise to streamline compound prioritization and facilitate drug repurposing, bridging the gap between computational inference and biological validation.

### Knowledge enhancement as an orthogonal and cost-efficient paradigm

Despite the rapid expansion of perturbation atlases and the emergence of single-cell foundation models, data-driven approaches for cellular perturbation modeling still face severe challenges, including the combinatorial complexity of the perturbation space, heterogeneous cellular contexts, and prohibitive experimental costs. Our findings introduce a crucial pivot: leveraging established biological knowledge as a strong, cost-efficient prior. We observe a clear, monotonic improvement in zero-shot performance as the scale of the knowledge graph expands (Section 2.5). And diverse biological knowledge, ranging from molecular structures to pathway interactions, synergistically contributes to the model’s generalization performance. This trend implies that mining broader and richer biological priors, such as unstructured scientific literature, offers a promising avenue for further improvement. Crucially, our paradigm is orthogonal to the underlying cell-state foundation model. Therefore, it provides a novel dimension that can be seamlessly combined with, rather than replaced by, data and model scaling to achieve systematic improvements.

### Future works

We identify two promising avenues to further advance this research. *First*, while this study focuses on small-molecule drugs, the proposed knowledge-driven pretraining naturally extends to a broader perturbation landscape, since it captures general biological priors between genes, chemical compounds and pathways. Future work could adapt our paradigm to model genetic interventions (*e*.*g*., CRISPR-mediated knockouts and knockdowns) or combinatorial treatments, as relevant datasets become available. *Second*, the rapid evolution of large-scale cell state foundation models offers a powerful basis for modeling cellular heterogeneity. Despite being computationally demanding, it’s a compelling direction to integrate our mechanistic knowledge directly into these foundation models.

## 4 Methods

MAP is a knowledge-enhanced framework for predicting cellular transcriptional responses to small-molecule perturbations. It addresses two key challenges: (i) generalizing to unseen cell type–drug combinations, and (ii) zero-shot prediction for drugs absent from training data. The core idea is to leverage structured biomedical knowledge to learn drug and gene representations that capture mechanistic similarities, enabling robust generalization across compounds and cellular contexts.

### 4.1 Problem Formulation

Given a small-molecule perturbation *p* ∈ 𝒫, our goal is to predict the cellular transcriptional response under a specific cell context. Formally, let *x* ∈ ℝ^*G*^ denote the gene expression profile of a cell over *G* genes under the control condition. Within a fixed cell context, we aim to learn a function that predicts the corresponding post-perturbation expression 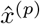:

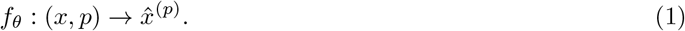

In practice, to reduce measurement noise, we perform *bulk-of-cells* prediction (*B* unperturbed cells *X* ∈ ℝ^*B×G*^) and take the average.

We evaluate generalization in two out-of-distribution (OOD) settings critical for drug discovery: **(i) novel cell type–drug combinations:** predicting responses for unseen combinations where both cell type and drug appear in training, but not together, addressing the sparsity of high-throughput screens; **(ii) unprofiled drugs:** predicting responses for compounds entirely absent from the training data (zero-shot learning), a capability essential for virtual screening of novel chemical entities.

### 4.2 Knowledge Graph Construction

We constructed **MAP-KG**, a large-scale biomedical knowledge graph that integrates structured and unstructured information spanning small-molecule drugs, genes, and their mechanistic relationships. MAP-KG aggregates curated evidence from 14 public databases (Supplementary Materials Section A.4), encompassing drug–target interactions, gene functional annotations, pathway memberships, and molecular mechanisms. By explicitly encoding biologically meaningful priors, MAP-KG provides a principled substrate for mechanism-aware modeling of cellular perturbation responses. As illustrated in Figure 1a, the graph comprises two core entity types—drugs and genes—and two classes of relational edges: drug–gene and gene–gene interactions. We describe each component below and provide more representative examples in Supplementary Table S14.

#### Drug nodes

Each drug node corresponds to a small-molecule compound and is uniquely indexed using a PubChem Compound Identifier (PubChem ID) [35]. We selected PubChem for its extensive coverage of chemical space and robust cross-referencing with pharmacological and molecular biology resources. In addition to the PubChem ID, each drug is associated with two attribute types: (i) **molecular structure (mol)**, encoded as SMILES strings; and (ii) **free form textual knowledge (text)**, comprising the compound name, descriptions of mechanisms of action (MOA) and biological therapeutic functions. Importantly, the MOA descriptions frequently reference target genes or proteins, thereby providing high-value semantic signals for modeling drug–gene relationships.

#### Gene nodes

Each gene node represents a human gene and is uniquely indexed by an Ensembl Gene Identifier (Ensembl ID) [36]. Ensembl IDs were chosen for their comprehensive coverage and stable, well-maintained mappings to alternative nomenclatures (*e*.*g*., HGNC [37]). Each gene node is further associated with two attribute types: (i) **the amino acid sequence (seq)**; (ii) **free form textual knowledge (text)**, spanning the gene symbol, molecular or cellular functional annotations, concise gene summaries, and descriptions of participating biological pathways. Together, these attributes capture complementary structural, functional, and contextual information about each gene.

#### Relation edges

Edges in MAP-KG encode biologically meaningful relationships between entities. Rather than constraining relations to a fixed ontology, we annotate each edge with a natural-language textual description, preserving semantic nuance and enabling flexible representation learning. The graph includes two principal categories of relations: **(i) drug–gene relations**, which capture mechanistic interactions such as binding, inhibition, activation, or transcriptional regulation. Each edge is defined by a PubChem ID for the drug node, an Ensembl ID for the gene node, a free-text description of the interaction, and an optional quantitative score reflecting interaction strength or confidence; **(ii) gene–gene relations**, which represent functional or physical associations, including regulatory interactions and protein–protein interactions. Similarly, each edge is identified by a pair of Ensembl IDs and annotated with descriptive text and optional quantitative scores. All edges in MAP-KG are **directed**, reflecting the asymmetric nature of biological relationships (e.g., regulator and target).

After normalization and deduplication, MAP-KG comprises 187,089 drug nodes, 22,924 gene nodes, 428,192 drug–gene edges, and 266,054 gene–gene edges. This graph offers a compact yet expressive representation of drug mechanisms and gene interactions, forming a biologically grounded foundation for learning knowledge-enhanced representations of cellular perturbations. Unlike previous efforts [21, 34, 38, 39], MAP-KG focuses on mechanistic knowledge linking genes and drugs: it integrates multimodal evidence and bridges modalities through rich textual descriptions, while also assembling a large-scale knowledge graph from diverse knowledge sources.

### 4.3 Knowledge-driven Multimodal Pre-training

To learn mechanism-aware representations, we pre-train the encoders on MAP-KG by aligning heterogeneous entity attributes and relational context in a unified embedding space (Figure 6).

**Figure 6.**
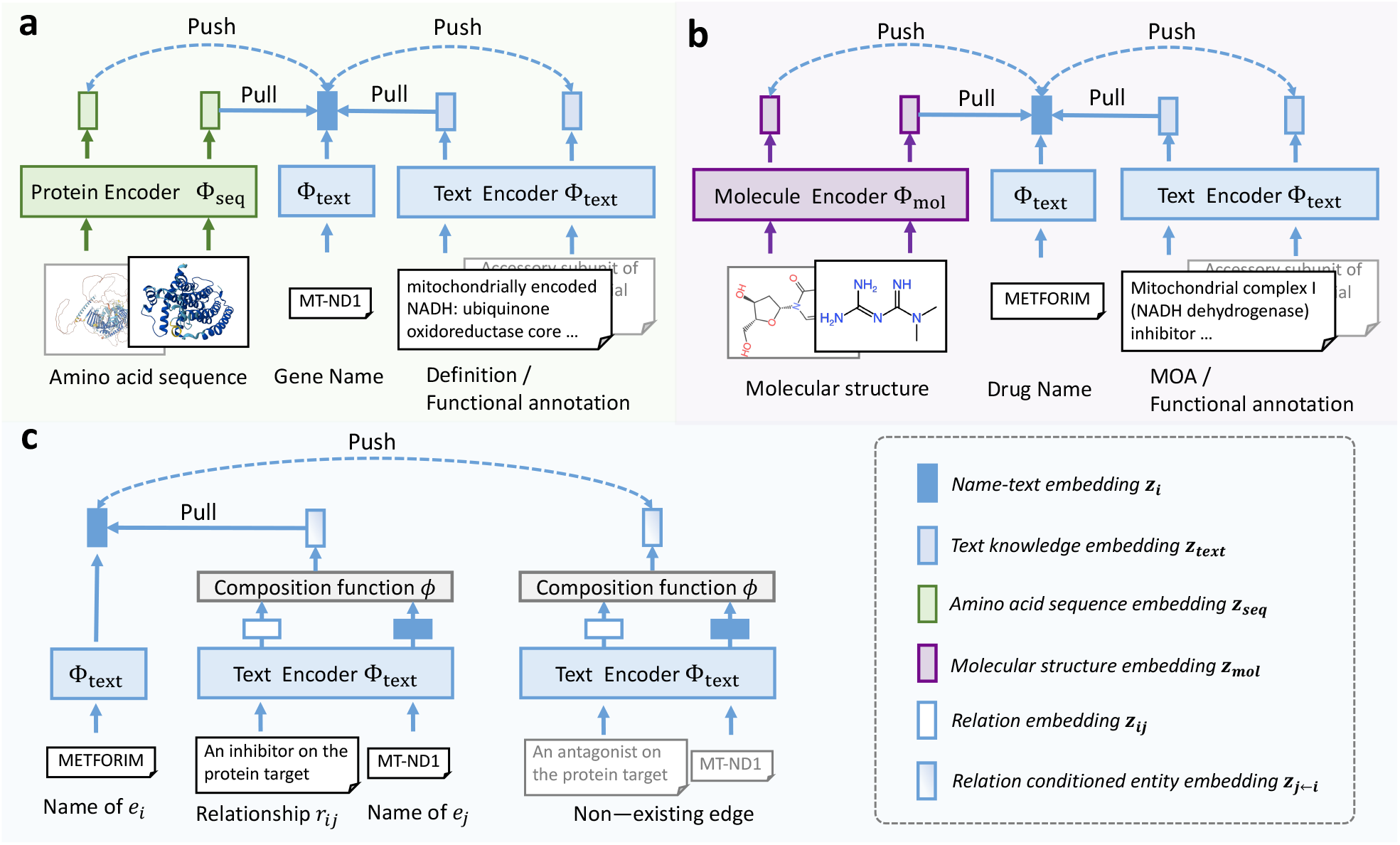
Knowledge-driven multimodal pre-training. We use modality-specific encoders to map heterogeneous biological input into the same vector space, and perform contrastive pre-training to align them, according to the biological priors. **(a)** and **(b)** show the inter-node attribute-level knowledge encoding and alignment for drug and gene entities. **(c)** shows the intra-node relation-level knowledge encoding and alignment between entities.

#### Intra-node knowledge encoding

As described in Section 4.2, each entity in MAP-KG is associated with intra-node knowledge from multiple modalities: (i) **text**, including entity names and natural-language descriptions (*e*.*g*., mechanism-of-action and functional annotations); (ii) **mol**, the molecular structure of drug nodes represented as SMILES strings; and (iii) **seq**, the canonical amino-acid sequence of gene nodes.

For each modality, we adopt a modality-specific encoder (Φ_text_, Φ_mol_, Φ_seq_) for knowledge encoding, where Φ_text_ is a biomedical language model (BioBERT [40]), Φ_mol_ is a molecule encoder for SMILES strings based on MoleculeSTM [33], and Φ_seq_ is a protein language model based on ESM-2 [41]. Specifically, we freeze the weights of ESM-2 and MoleculeSTM to prevent feature degradation, and append 4 trainable MLP layers to each of them for feature adaptation. Let ℰ denote the set of all entities (drugs and genes) in the graph, and each entity *e*_*i*_ ∈ ℰ, denotes one of its modality-specific attributes as *k*_*i*_ ∈ {*k*_text_, *k*_mol_, *k*_seq_}:

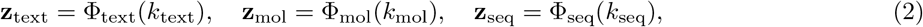

where the encoders project heterogeneous biological inputs into a *d*-dimensional vector, which will be aligned later via contrastive pre-training. For simplicity, we use the name-text (*i*.*e*., compound name or gene symbol) embedding from Φ_text_ as the *canonical representation* for *e*_*i*_, denoted as **z**_*i*_.

#### Inter-node relation encoding

Inter-node relational knowledge in MAP-KG is expressed as natural-language descriptions on directed edges. For an edge from *e*_*i*_ to *e*_*j*_ with relation text *r*_*ij*_ (e.g., “inhibits”), we encode the relation using the text encoder:

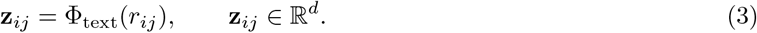

To explicitly model how an entity participates in a specific relation, we construct **relation-conditioned entity embeddings**. Specifically, we introduce a relation-conditioned composition function *ϕ*_rc_(·), implemented as a multi-layer perceptron (MLP), which maps an *ordered* pair of embeddings to a *d*-dimensional representation:

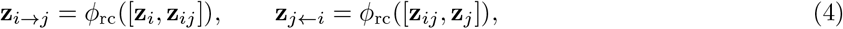

where [·,·] denote concatenation, **z**_*i*_ and **z**_*j*_ are canonical representations of *e*_*i*_ and *e*_*j*_, **z**_*i*→*j*_ denotes the embedding of entity *e*_*i*_ conditioned on its relation to *e*_*j*_, and **z**_*j*←*i*_ denotes the embedding of *e*_*j*_ conditioned on the relation from *e*_*i*_. This order-sensitive composition function enables the model to distinguish the source entity from the target entity in directed relations. For instance, given the triplet (*Metformin, an inhibitor on the protein target, MT-ND1*) in Figure 1(a), **z**_*i*→*j*_ encodes Metformin’s active role as the regulator exerting inhibition, whereas **z**_*j*←*i*_ characterizes MT-ND1 in the passive state of being targeted. This design captures the inherent directionality of biological interactions, effectively modeling the causal roles encoded in the graph edges.

#### Contrastive pre-training

As shown in Figure 6, we train the knowledge encoders with contrastive learning. Given a mini-batch ℬ, for each entity *e*_*i*_, we first construct a pair of embeddings 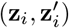 that encode a specific biological prior and should be aligned in the representation geometry. Here, **z**_*i*_ is the canonical representation, *i*.*e*., the name-text embedding of *e*_*i*_, and 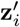 is randomly sampled according to two supervision strategies: **(i) intra-node attribute-level knowledge alignment** (Figure 6a) pulls the attributes of each entity towards its name in the embedding space. In this case, 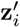 is instantiated from { **z**_text_, **z**_mol_, **z**_seq_ }, derived in Equation 2; **(ii) inter-node relation-level knowledge alignment** (Figure 6b) pulls the relation-conditioned entity towards the linked neighbor in the embedding space. Here, 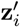 is instantiated as **z**_*j←i*_, given a directed relationship from entity *e*_*j*_, or **z**_*j*_→_*i*_, given a directed relationship towards entity *e*_*j*_, following Equation 4. Then, we minimize the InfoNCE loss [42] over the mini-batch:

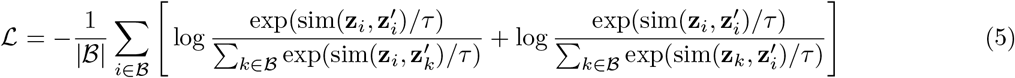

where *τ* is a temperature hyperparameter.

#### Discussion

The proposed knowledge-driven multi-modal pre-training embeds drugs and genes in a shared space by aligning heterogeneous and fragmented biomedical evidence, so that mechanistically or functionally similar drugs form coherent neighborhoods in the representation geometry, rather than being isolated. Consequently, the geometry provides a transferable inductive bias: even when downstream perturbation data are sparse and noisy, the model can generalize by propagating information through these similarity-aware embeddings.

### 4.4 Knowledge-guided perturbation response prediction

In this section, we describe how knowledge-informed drug and gene representations are incorporated with a pretrained single-cell foundation model to guide perturbation response prediction (Figure 7). Formally, given a bulk *X* ∈ ℝ^*B×G*^ of *B* cells under the control condition, for each cell *x* ∈ ℝ^*G*^ in the bulk, we first encode the unperturbed cellular state using a pretrained foundation model (STATE SE-600M) [15]:

**Figure 7.**
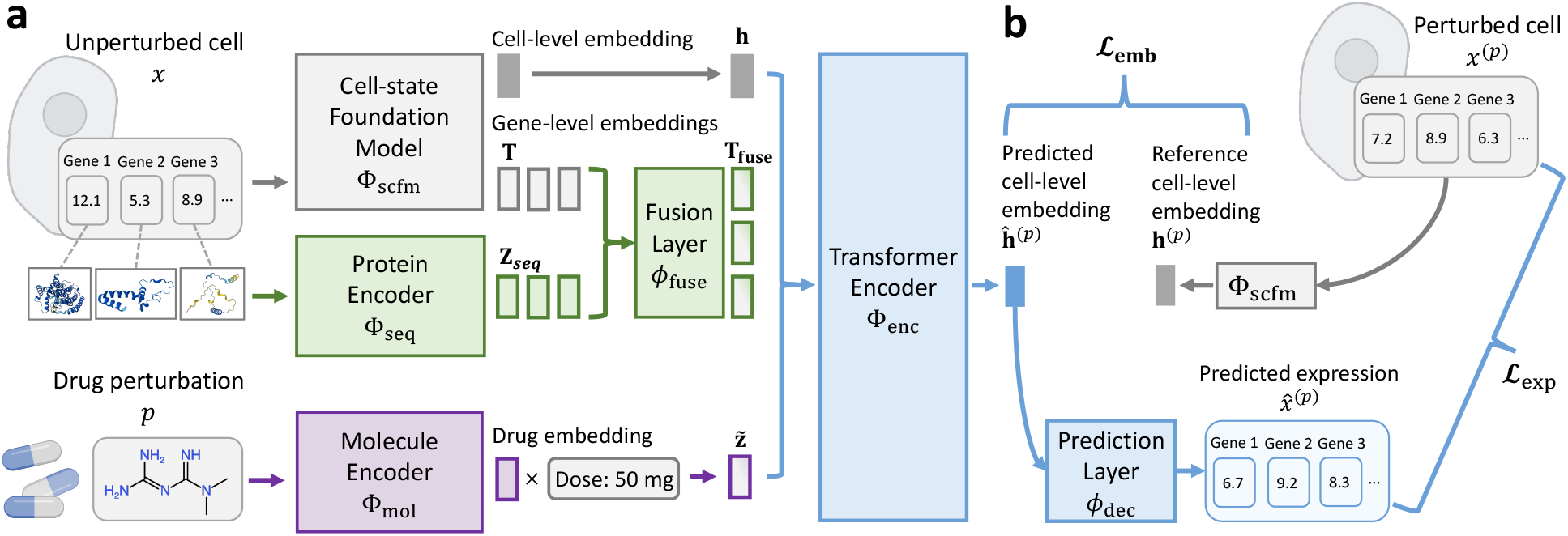
Knowledge-guided perturbation response prediction. **(a)** We devise a framework that seamlessly integrates the knowledge-informed drug and gene representations with a cell-state foundation model, to infer the perturbation response. **(b)** It is trained with both expression-level and embedding-level supervision to alleviate the measurement noise.

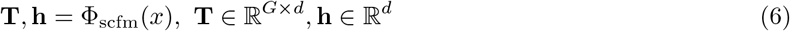

where **T** are the latent gene-level embeddings, and **h** is the cell-level embedding that captures the global unperturbed cellular state, taken as the output of the prepended `SPECIAL` token. Then, we enhance the gene embeddings **T** with knowledge-informed gene-level representations **z**_seq_ derived in Section 4.3, and fuse them via a multi-layer perceptron *ϕ*_fuse_:

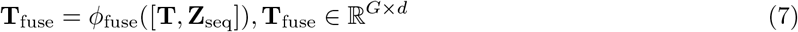

where **Z**_seq_ ∈ ℝ^*G×d*^ is obtained by stacking the knowledge-informed representations of all *G* genes, and [·, ·] denotes concatenation along the feature dimension.

For a given perturbation *p* ∈ 𝒫, we take its molecular SMILES string, obtain the knowledge-informed representation **z**_mol_ using the pre-trained Φ_mol_ described in Section 4.3, and scale the representation by 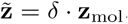, where *δ* ∈ *R* is the log-transformed dose value applied. A transformer encoder module with 4 self-attention layers is then adopted to infer the perturbed cell-level embeddings. It takes as input a sequence constructed by concatenating 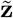 and **T**_fuse_:

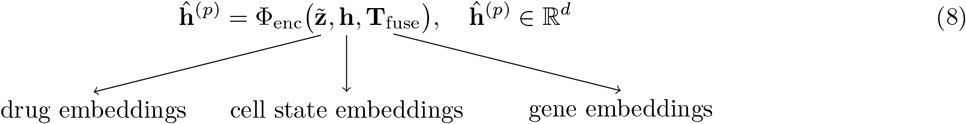

Here, **ĥ**^(*p*)^ corresponds to the output embedding of **h**. The perturbed cell embeddings are then passed through a multi-layer perceptron head *ϕ*_dec_ to decode the perturbed gene expression profile:

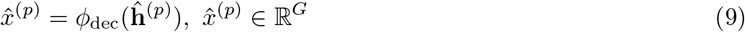

For a bulk *X*, the above procedure is applied independently to each cell. We denote by 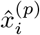 and 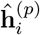 the predicted expression and embedding for the *i*-th cell under perturbation *p*. The bulk-level predictions 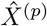 and **Ĥ** ^(*p*)^ are obtained by stacking these cell-wise outputs.

#### Training objective

Following previous works [15], we train the perturbation predictor using a dual-space supervision objective that matches both (i) the perturbed expression profile in data space and (ii) the perturbed cellular embedding in the STATE latent space. Specifically, for each training tuple (*X, p, X*^(*p*)^), we first obtain the reference embedding matrix **H**^(*p*)^ by encoding each authentic perturbed cell 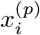 with the foundation model Φ_scfm_ and stacking the resulting cell-level embeddings 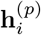.

The loss function is then formulated as:

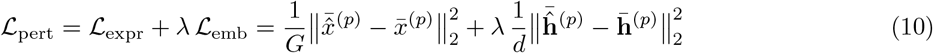

where *λ* balances the two terms, and 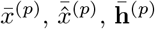 and 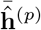 denote the cell-wise means within the bulk:

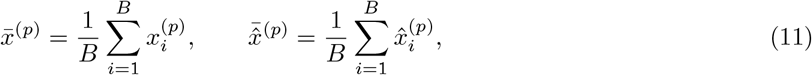

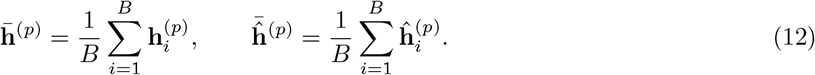

Here, the embedding-level supervision distills the perturbation-induced shift captured by the foundation model, providing a stable training signal when expression-level supervision is noisy or sparse.

#### Discussion

Since each drug is represented by its SMILES string, the model is naturally extensible to any novel compounds at inference time in a zero-shot manner, as long as their SMILES strings are available. Moreover, Φ_scfm_ can be seamlessly replaced by other pre-trained single-cell foundation models, making the proposed method broadly applicable across different backbone choices. Put differently, improvements from scaling data and single-cell foundation model are orthogonal to our contributions.

### 4.5 Datasets

We evaluated MAP on three large-scale single-cell perturbation datasets that profile transcriptional responses to small-molecule drug treatments across diverse cellular contexts. All datasets consist of paired control and perturbed single-cell RNA-sequencing (scRNA-seq) profiles and were processed using a unified preprocessing pipeline to ensure consistency across experiments. Across all datasets, vehicle-only conditions were treated as controls (DMSO when available; otherwise the dataset-provided vehicle), and all compound-treated conditions were regarded as perturbations.

#### 4.5.1 Datasets descriptions

**Tahoe-100M dataset** [23] is a giga-scale single-cell perturbation atlas spanning cancer models. It comprises 95.6 million quality-controlled scRNA-seq profiles collected from 50 cancer cell lines, of which 47 were retained for downstream analysis due to sufficient coverage. The dataset includes 379 drugs and 1,135 drug–dose combinations, yielding 52,886 perturbation conditions defined by the combination of cell line, compound, and dose (median 1,287 cells per condition). To provide a proof-of-concept evaluation while maintaining computational tractability, we restricted training and validation to six cell lines spanning diverse tissues and cancer types. For each cell line, highly variable genes (HVGs) were identified based on within–cell-line expression variability. To evaluate zero-shot generalization to unseen cell line–drug combinations, we held out 5% of drugs for each cell line as test drugs, with non-overlapping test sets across cell lines. To assess zero-shot generalization to entirely unprofiled drugs, we additionally held out 5% of all drugs (*n* = 16) during training.

**OP3 dataset** [24] is a Peripheral Blood Mononuclear Cells (PBMCs)-based single-cell perturbation benchmark. It contains 240,059 cells with expression measured over 18,211 genes under 147 perturbation conditions, including 144 compounds administered at 1 µM. DMSO at 14.1 µM serves as the negative control, and two compounds (dabrafenib and belinostat at 0.1 µM) are included as positive controls. Cells are annotated into six immune cell types (CD4^+^ T, CD8^+^ T, NK, regulatory T, B, and myeloid populations); for B cells and myeloid cells, only a subset of drug conditions is available. All six cell types were included in our experiments. Following the same splitting strategy as for Tahoe-100M, we held out 5% of drugs per cell type to evaluate generalization to unseen cell type–drug combinations, and an additional 5% of all drugs to assess generalization to unseen drugs.

**SciPlex3 dataset** [2] is a single-cell chemical perturbation dataset spanning three human cancer cell lines: A549 (lung adenocarcinoma), MCF7 (breast adenocarcinoma), and K562 (chronic myelogenous leukemia). It contains 581,777 cells treated with 187 compounds, with expression quantified over 58,347 genes. All three cell lines were included in our experiments, and we adopted the same train–test splitting scheme as described for Tahoe-100M.

#### 4.5.2 Data preprocessing

##### Gene expression preprocessing

For single-cell foundation model inputs, we restricted the gene space to 19,790 human protein-coding genes defined by Ensembl annotations. Each cell was normalized to a total of 10,000 unique molecular identifier (UMI) counts and log-transformed via log 1*p* to obtain normalized expression values. For each cell, we selected the top *L* = 2,048 genes ranked by log-normalized expression to form a fixed-length gene token sequence as input to the foundation model. Expression magnitudes were encoded using the soft-binning strategy described in [15]. This preprocessing pipeline was applied consistently across all three datasets.

##### Molecular structure preprocessing

For each compound, we used RDKit [43] to standardize its SMILES string and generate the canonical SMILES representation, which served as input to the molecule encoder.

##### Prediction targets

We evaluated predictions on two complementary gene sets: HVGs and DEGs. Highly variable genes (HVGs), defined as genes exhibiting the largest expression variability across cells and capturing informative biological heterogeneity, were identified directly from each dataset’s `AnnData` object using Scanpy [44]. Across all experiments, we consistently used the top 2,000 HVGs as prediction targets. Differentially expressed genes (DEGs), defined as genes exhibiting significant expression changes between perturbed and control cells for a given condition, were likewise identified using Scanpy. For both HVGs and DEGs, we applied library-size normalization followed by a log1p transformation.

## 5 Data Availability

We obtain the Tahoe-100M dataset from Huggingface, OP3 dataset from NeurIPS competition and SciPlex3 dataset from GEO. We release MAP-KG at https://huggingface.co/datasets/RainGate/MAP-KG, and provide the detailed knowledge source in Supplementary Table S13.

## 6 Code Availability

All the code and models are publicly accessible at https://github.com/MAGIC-AI4Med/MAP.

## A Supplementary Materials

### A.1 Detailed experimental results

This section provides the detailed numerical results corresponding to the key findings reported in the main-text Results section. The tables are arranged to follow the order of the Results section. Tables S1 to S3 show results of the zero-shot generalization to unseen cell type-drug combinations (Section 2.2). Tables S4 to S9 show results of the zero-shot generalization to unprofiled drugs (Section 2.3), Table S10 shows the GSEA results of the four prioritized anti-cancer drugs (Section 2.4), and Tables S11 and S12 show analysis of knowledge scale and knowledge type diversity (Section 2.5).

**Table S1.** Average performance comparison between MAP and baseline models on 6 cell lines selected from Tahoe-100M dataset, under the regime of zero-shot generalization to unseen cell type-drug combinations. Pearson delta correlation, Perturbation discrimination score and direction accuracy are calculated. 95% confidence intervals are reported.

| Method | HVG |  |  | Top-50 DEG |  |
| --- | --- | --- | --- | --- | --- |
| | Pearson $\Delta$ | PDS | Direction Acc | Pearson $\Delta$ | Direction Acc |
| **trainMean** | 0.331 ± 0.005 | 0.024 ± 0.001 | 0.553 ± 0.016 | 0.219 ± 0.004 | 0.741 ± 0.014 |
| **PRnet** | 0.327 ± 0.004 | 0.096 ± 0.003 | 0.662 ± 0.017 | 0.229 ± 0.004 | 0.618 ± 0.012 |
| **chemCPA** | 0.677 ± 0.014 | 0.285 ± 0.004 | 0.636 ± 0.015 | 0.724 ± 0.021 | 0.832 ± 0.013 |
| **CRISP** | 0.693 ± 0.015 | 0.754 ± 0.028 | 0.608 ± 0.005 | 0.797 ± 0.017 | 0.855 ± 0.022 |
| **MAP** | **0.878 ± 0.019** | **0.906 ± 0.013** | **0.791 ± 0.029** | **0.930 ± 0.005** | **0.990 ± 0.016** |

**Table S2.** Cell type specific analysis of Tahoe-100M dataset, under the regime of zero-shot generalization to unseen cell type-drug combinations. We report cell-line-wise performance (top-50 DEG Pearson delta correlation and direction accuracy) on 6 curated cell lines, averaged across testing drug sets of each cell line.

| Model | A-172 | A-498 | A-549 | Hep-G2/C3A | PANC-1 | SK-MEL-2 | Average |
| --- | --- | --- | --- | --- | --- | --- | --- |
| *Metric: Pearson $\Delta$ top-50 DEG* |  |  |  |  |  |  |  |
| **trainMean** | 0.330 ± 0.014 | 0.486 ± 0.015 | 0.338 ± 0.013 | 0.347 ± 0.019 | 0.514 ± 0.013 | 0.521 ± 0.024 | 0.423 ± 0.016 |
| **PRnet** | 0.380 ± 0.003 | 0.590 ± 0.004 | 0.363 ± 0.023 | 0.462 ± 0.003 | 0.572 ± 0.003 | 0.497 ± 0.024 | 0.477 ± 0.010 |
| **chemCPA** | 0.768 ± 0.020 | 0.682 ± 0.034 | 0.761 ± 0.019 | 0.717 ± 0.021 | 0.780 ± 0.018 | 0.733 ± 0.015 | 0.740 ± 0.021 |
| **CRISP** | 0.750 ± 0.009 | 0.837 ± 0.021 | 0.831 ± 0.018 | 0.764 ± 0.020 | 0.801 ± 0.027 | 0.793 ± 0.013 | 0.795 ± 0.018 |
| **MAP** | **0.957 ± 0.007** | **0.945 ± 0.006** | **0.915 ± 0.007** | **0.895 ± 0.011** | **0.935 ± 0.017** | **0.925 ± 0.009** | **0.929 ± 0.009** |
| *Metric: Direction Acc top-50 DEG* |  |  |  |  |  |  |  |
| **trainMean** | 0.670 ± 0.011 | 0.841 ± 0.019 | 0.682 ± 0.016 | 0.720 ± 0.019 | 0.808 ± 0.015 | 0.686 ± 0.007 | 0.734 ± 0.014 |
| **PRnet** | 0.553 ± 0.012 | 0.678 ± 0.012 | 0.503 ± 0.010 | 0.584 ± 0.010 | 0.688 ± 0.013 | 0.700 ± 0.018 | 0.617 ± 0.012 |
| **chemCPA** | 0.804 ± 0.017 | 0.872 ± 0.034 | 0.840 ± 0.015 | 0.809 ± 0.010 | 0.851 ± 0.019 | 0.838 ± 0.016 | 0.835 ± 0.018 |
| **CRISP** | 0.822 ± 0.020 | 0.908 ± 0.021 | 0.869 ± 0.015 | 0.832 ± 0.034 | 0.879 ± 0.021 | 0.868 ± 0.022 | 0.863 ± 0.022 |
| **MAP** | **0.995 ± 0.005** | **0.992 ± 0.006** | **0.985 ± 0.012** | **0.978 ± 0.020** | **0.990 ± 0.008** | **0.988 ± 0.010** | **0.988 ± 0.010** |

**Table S3.** Cell type specific analysis of OP3 and SciPlex3 datasets, under the regime of zero-shot generalization to unseen cell type-drug combinations. We show fine-grained performance (top-50 DEG Pearson delta correlation) on 6 cell types of OP3 (B cells, Myeloid cells, NK cells, T cells CD4+, T cells CD8+ and T regulatory cells) and 3 cell lines of SciPlex3 (A-549, K-562 and MCF-7, averaged across testing drug sets of each cell type.

|  | MAP | CRISP | chemCPA | PRnet | trainMean |
| --- | --- | --- | --- | --- | --- |
| **B cells** | **0.815 ± 0.017** | 0.714 ± 0.011 | 0.690 ± 0.010 | 0.556 ± 0.023 | 0.594 ± 0.015 |
| **Myeloid cells** | **0.642 ± 0.007** | 0.624 ± 0.021 | 0.594 ± 0.013 | 0.492 ± 0.035 | 0.542 ± 0.015 |
| **NK cells** | **0.693 ± 0.015** | 0.673 ± 0.014 | 0.648 ± 0.012 | 0.504 ± 0.027 | 0.560 ± 0.015 |
| **T cells CD4+** | **0.713 ± 0.011** | 0.684 ± 0.012 | 0.671 ± 0.013 | 0.516 ± 0.031 | 0.570 ± 0.012 |
| **T cells CD8+** | **0.689 ± 0.008** | 0.653 ± 0.011 | 0.639 ± 0.005 | 0.506 ± 0.023 | 0.546 ± 0.016 |
| **T regulatory cells** | **0.724 ± 0.012** | 0.697 ± 0.022 | 0.675 ± 0.014 | 0.524 ± 0.034 | 0.579 ± 0.010 |
| **Average** | **0.706 ± 0.014** | 0.673 ± 0.009 | 0.650 ± 0.013 | 0.524 ± 0.048 | 0.568 ± 0.018 |
| **A-549** | **0.808 ± 0.009** | 0.764 ± 0.018 | 0.617 ± 0.023 | 0.503 ± 0.024 | 0.456 ± 0.006 |
| **K-562** | **0.857 ± 0.006** | 0.812 ± 0.011 | 0.783 ± 0.010 | 0.574 ± 0.012 | 0.402 ± 0.004 |
| **MCF7** | **0.769 ± 0.008** | 0.615 ± 0.023 | 0.682 ± 0.009 | 0.461 ± 0.013 | 0.441 ± 0.006 |
| **Average** | **0.811 ± 0.008** | 0.724 ± 0.015 | 0.697 ± 0.015 | 0.512 ± 0.018 | 0.433 ± 0.006 |

**Table S4.** Performance comparison between MAP and baseline models on 6 cell lines selected from Tahoe-100M dataset, perturbed with seen drugs. 95% confidence intervals are reported. Best results are highlighted in bold.

| Metric | MAP | STATE | CRISP | chemCPA | PRnet | trainMean |
| --- | --- | --- | --- | --- | --- | --- |
| Pearson $\Delta$ HVG | **0.869 ± 0.023** | 0.827 ± 0.023 | 0.721 ± 0.019 | 0.718 ± 0.018 | 0.262 ± 0.007 | 0.331 ± 0.009 |
| PDS HVG | **0.992 ± 0.026** | 0.911 ± 0.018 | 0.790 ± 0.016 | 0.266 ± 0.006 | 0.057 ± 0.001 | 0.017 ± 0.001 |
| Direction Acc HVG | **0.816 ± 0.019** | 0.731 ± 0.019 | 0.710 ± 0.017 | 0.664 ± 0.017 | 0.637 ± 0.019 | 0.454 ± 0.006 |
| Pearson $\Delta$ top-50 DEG | **0.969 ± 0.021** | 0.943 ± 0.016 | 0.896 ± 0.023 | 0.711 ± 0.014 | 0.312 ± 0.010 | 0.461 ± 0.016 |
| Direction Acc top-50 DEG | **0.987 ± 0.013** | 0.922 ± 0.026 | 0.842 ± 0.026 | 0.846 ± 0.015 | 0.565 ± 0.011 | 0.689 ± 0.020 |

**Table S5.** Performance comparison between MAP and baseline models on 6 cell lines selected from Tahoe-100M dataset, perturbed with unseen drugs. 95% confidence intervals are reported. Best results are highlighted in bold.

| Metric | MAP | CRISP | chemCPA | PRnet | trainMean |
| --- | --- | --- | --- | --- | --- |
| Pearson $\Delta$ HVG | **0.659 ± 0.014** | 0.489 ± 0.012 | 0.202 ± 0.004 | 0.223 ± 0.005 | 0.277 ± 0.005 |
| PDS HVG | **0.344 ± 0.006** | 0.175 ± 0.005 | 0.022 ± 0.000 | 0.003 ± 0.000 | 0.013 ± 0.000 |
| Direction Acc HVG | **0.807 ± 0.021** | 0.655 ± 0.021 | 0.567 ± 0.009 | 0.632 ± 0.013 | 0.470 ± 0.009 |
| Pearson $\Delta$ top-50 DEG | **0.822 ± 0.023** | 0.700 ± 0.016 | 0.671 ± 0.015 | 0.327 ± 0.009 | 0.428 ± 0.010 |
| Direction Acc top-50 DEG | **0.934 ± 0.024** | 0.724 ± 0.020 | 0.559 ± 0.012 | 0.531 ± 0.014 | 0.693 ± 0.013 |

**Table S6.** Compound-specific analysis under the regime of zero-shot generalization to unprofiled drugs. Top 50 DE genes’ Pearson delta correlation are reported with the 95% confidence intervals.

| Compound | MAP | CRISP | chemCPA | PRnet | trainMean |
| --- | --- | --- | --- | --- | --- |
| Balsalazide (sodium hydrate) | **0.854 ± 0.029** | 0.836 ± 0.068 | 0.575 ± 0.043 | 0.178 ± 0.486 | 0.602 ± 0.088 |
| Bergenin | **0.802 ± 0.076** | 0.504 ± 0.038 | 0.556 ± 0.047 | 0.140 ± 0.251 | 0.203 ± 0.094 |
| Brivudine | **0.678 ± 0.145** | 0.555 ± 0.045 | 0.594 ± 0.049 | 0.282 ± 0.206 | 0.559 ± 0.126 |
| Carbidopa (monohydrate) | **0.965 ± 0.033** | 0.884 ± 0.068 | 0.763 ± 0.052 | 0.425 ± 0.102 | 0.382 ± 0.103 |
| Ciclopirox | **0.620 ± 0.042** | 0.541 ± 0.034 | 0.532 ± 0.041 | 0.222 ± 0.224 | 0.313 ± 0.104 |
| CP21R7 | **0.865 ± 0.043** | 0.652 ± 0.049 | 0.432 ± 0.033 | 0.130 ± 0.229 | 0.424 ± 0.108 |
| Drospirenone | **0.941 ± 0.058** | 0.882 ± 0.059 | 0.745 ± 0.053 | 0.487 ± 0.316 | 0.242 ± 0.130 |
| ERK5-IN-2 | **0.869 ± 0.041** | 0.635 ± 0.051 | 0.743 ± 0.055 | 0.324 ± 0.219 | 0.419 ± 0.086 |
| Estrone sulfate (potassium) | **0.816 ± 0.064** | 0.706 ± 0.050 | 0.653 ± 0.026 | 0.346 ± 0.072 | 0.260 ± 0.064 |
| Idarubicin (hydrochloride) | **0.737 ± 0.072** | 0.458 ± 0.035 | 0.582 ± 0.043 | 0.385 ± 0.263 | 0.297 ± 0.091 |
| Lidocaine (hydrochloride) | **0.932 ± 0.034** | 0.690 ± 0.028 | 0.733 ± 0.057 | 0.329 ± 0.218 | 0.282 ± 0.068 |
| Nafamostat (mesylate) | **0.808 ± 0.193** | 0.701 ± 0.055 | 0.700 ± 0.056 | 0.497 ± 0.080 | 0.365 ± 0.072 |
| Ralimetinib dimesylate | **0.757 ± 0.027** | 0.623 ± 0.049 | 0.680 ± 0.057 | 0.612 ± 0.228 | 0.421 ± 0.072 |
| Sildenafil | **0.942 ± 0.038** | 0.852 ± 0.068 | 0.775 ± 0.055 | 0.597 ± 0.236 | 0.412 ± 0.050 |
| Sivelestat (sodium tetrahydrate) | **0.916 ± 0.022** | 0.837 ± 0.059 | 0.789 ± 0.063 | 0.559 ± 0.240 | 0.408 ± 0.046 |
| ULK-101 | **0.713 ± 0.095** | 0.610 ± 0.054 | 0.572 ± 0.035 | 0.344 ± 0.221 | 0.364 ± 0.045 |

**Table S7.** Compound-specific analysis under the regime of zero-shot generalization to unprofiled drugs. Top 50 DE genes’ direction accuracy are reported with the 95% confidence intervals.

| Compound | MAP | CRISP | chemCPA | PRnet | trainMean |
| --- | --- | --- | --- | --- | --- |
| Balsalazide (sodium hydrate) | **0.937 ± 0.025** | 0.605 ± 0.022 | 0.676 ± 0.047 | 0.700 ± 0.039 | 0.815 ± 0.039 |
| Bergenin | **0.930 ± 0.049** | 0.876 ± 0.035 | 0.737 ± 0.046 | 0.526 ± 0.028 | 0.616 ± 0.088 |
| Brivudine | **0.841 ± 0.066** | 0.839 ± 0.062 | 0.820 ± 0.070 | 0.622 ± 0.043 | 0.868 ± 0.090 |
| Carbidopa (monohydrate) | **0.959 ± 0.025** | 0.728 ± 0.054 | 0.715 ± 0.052 | 0.468 ± 0.029 | 0.673 ± 0.070 |
| Ciclopirox | **0.862 ± 0.065** | 0.636 ± 0.044 | 0.390 ± 0.035 | 0.524 ± 0.033 | 0.739 ± 0.065 |
| CP21R7 | **0.938 ± 0.048** | 0.851 ± 0.062 | 0.762 ± 0.072 | 0.582 ± 0.027 | 0.680 ± 0.050 |
| Drospirenone | **0.968 ± 0.029** | 0.688 ± 0.056 | 0.626 ± 0.046 | 0.475 ± 0.070 | 0.533 ± 0.114 |
| ERK5-IN-2 | **0.928 ± 0.034** | 0.895 ± 0.065 | 0.544 ± 0.041 | 0.595 ± 0.037 | 0.572 ± 0.069 |
| Estrone sulfate (potassium) | **0.971 ± 0.032** | 0.783 ± 0.063 | 0.720 ± 0.068 | 0.495 ± 0.023 | 0.744 ± 0.040 |
| Idarubicin (hydrochloride) | **0.944 ± 0.021** | 0.782 ± 0.052 | 0.441 ± 0.031 | 0.567 ± 0.027 | 0.421 ± 0.048 |
| Lidocaine (hydrochloride) | **0.970 ± 0.028** | 0.736 ± 0.058 | 0.708 ± 0.045 | 0.527 ± 0.023 | 0.732 ± 0.062 |
| Nafamostat (mesylate) | **0.826 ± 0.051** | 0.445 ± 0.030 | 0.758 ± 0.028 | 0.533 ± 0.039 | 0.676 ± 0.078 |
| Ralimetinib dimesylate | **0.889 ± 0.046** | 0.485 ± 0.031 | 0.682 ± 0.057 | 0.566 ± 0.056 | 0.735 ± 0.068 |
| Sildenafil | **0.985 ± 0.030** | 0.907 ± 0.049 | 0.782 ± 0.053 | 0.492 ± 0.082 | 0.606 ± 0.036 |
| Sivelestat (sodium tetrahydrate) | **0.969 ± 0.065** | 0.800 ± 0.031 | 0.693 ± 0.052 | 0.496 ± 0.026 | 0.736 ± 0.038 |
| ULK-101 | **0.896 ± 0.037** | 0.746 ± 0.069 | 0.539 ± 0.044 | 0.592 ± 0.075 | 0.821 ± 0.054 |

**Table S8.** Average performance comparison between MAP and baseline models on OP3 dataset, under the regime of zero-shot generalization to unprofiled drugs. Pearson delta correlation, perturbation discrimination score and direction accuracy are reported.

| Method | HVG |  |  | Top-50 DEG |  |
| --- | --- | --- | --- | --- | --- |
| | Pearson $\Delta$ | PDS | Direction Acc | Pearson $\Delta$ | Direction Acc |
| **trainMean** | $0.239 \pm 0.015$ | $0.016 \pm 0.001$ | $0.418 \pm 0.009$ | $0.468 \pm 0.009$ | $0.658 \pm 0.013$ |
| **PRnet** | $0.245 \pm 0.008$ | $0.048 \pm 0.003$ | $0.456 \pm 0.013$ | $0.415 \pm 0.012$ | $0.686 \pm 0.013$ |
| **chemCPA** | $0.473 \pm 0.008$ | $0.090 \pm 0.003$ | $0.536 \pm 0.014$ | $0.663 \pm 0.009$ | $0.783 \pm 0.014$ |
| **CRISP** | $0.525 \pm 0.012$ | $0.106 \pm 0.005$ | $0.574 \pm 0.017$ | $0.652 \pm 0.014$ | $0.815 \pm 0.020$ |
| **MAP** | **$0.601 \pm 0.021$** | **$0.155 \pm 0.009$** | **$0.692 \pm 0.019$** | **$0.734 \pm 0.007$** | **$0.921 \pm 0.014$** |

**Table S9.**
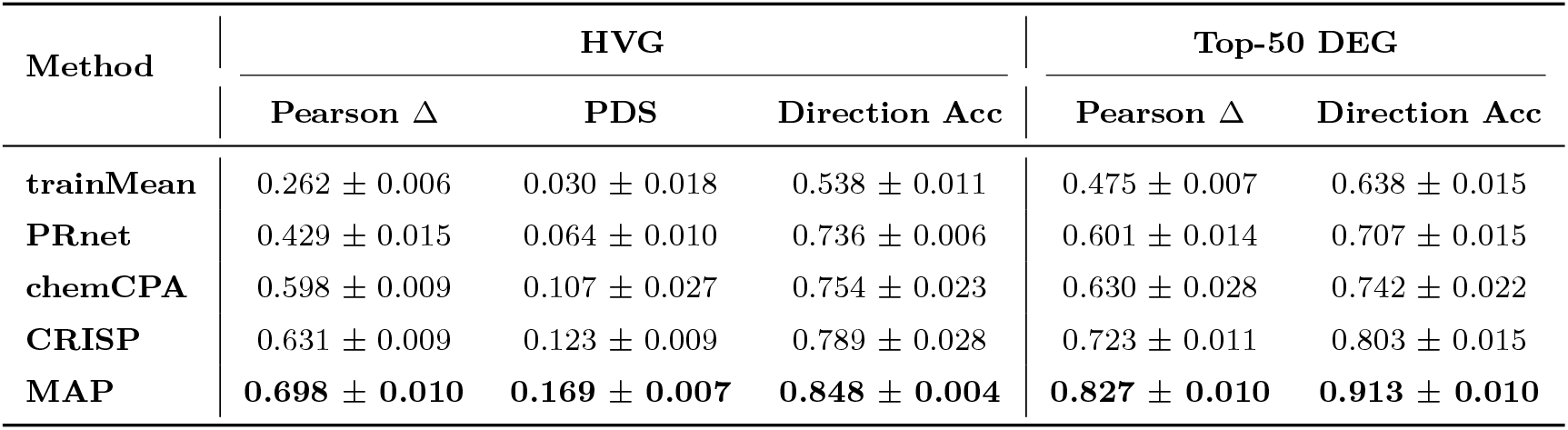
Average performance comparison between MAP and baseline models on SciPlex3 dataset, under the regime of zero-shot generalization to unprofiled drugs. Pearson delta correlation, perturbation discrimination score and direction accuracy are reported.

| Method | HVG |  |  | Top-50 DEG |  |
| --- | --- | --- | --- | --- | --- |
| | Pearson $\Delta$ | PDS | Direction Acc | Pearson $\Delta$ | Direction Acc |
| **trainMean** | $0.262 \pm 0.006$ | $0.030 \pm 0.018$ | $0.538 \pm 0.011$ | $0.475 \pm 0.007$ | $0.638 \pm 0.015$ |
| **PRnet** | $0.429 \pm 0.015$ | $0.064 \pm 0.010$ | $0.736 \pm 0.006$ | $0.601 \pm 0.014$ | $0.707 \pm 0.015$ |
| **chemCPA** | $0.598 \pm 0.009$ | $0.107 \pm 0.027$ | $0.754 \pm 0.023$ | $0.630 \pm 0.028$ | $0.742 \pm 0.022$ |
| **CRISP** | $0.631 \pm 0.009$ | $0.123 \pm 0.009$ | $0.789 \pm 0.028$ | $0.723 \pm 0.011$ | $0.803 \pm 0.015$ |
| **MAP** | **$0.698 \pm 0.010$** | **$0.169 \pm 0.007$** | **$0.848 \pm 0.004$** | **$0.827 \pm 0.010$** | **$0.913 \pm 0.010$** |

**Table S10.** GSEA Results for Four Drugs on A-549 Cell Line. Normalized Enrichment Scores (NES) and False Discovery Rate-adjusted q value (FDR-q) of predictions and values are reported for each gene set (pathway). For every drug, 12 representative gene sets that are linked to each compound’s known mechanism of action and to disease-relevant699 programs in A549

| Drug | Pathway Name | True NES | True FDR-q | Pred NES | Pred FDR-q |
| --- | --- | --- | --- | --- | --- |
| Afatinib | NF-κB Signaling | 1.991 | 0.199 | 1.451 | 0.210 |
|  | LPS Inflammatory Response | 1.913 | 0.031 | 1.175 | 0.041 |
|  | Apoptosis (WP) | 1.586 | 0.091 | 1.312 | 0.194 |
|  | MYC E-box Targets | 1.608 | 0.188 | 1.399 | 0.232 |
| | NF-κB Fibroblast Targets | 1.752 | 0.417 | 1.467 | 0.601 |
|  | EGF/EGFR Signaling (WP) | -0.980 | 0.120 | -1.294 | 0.190 |
|  | EGF Response (HeLa) | -0.827 | 0.125 | -0.766 | 0.230 |
|  | ERBB2 Isoform-1 Metastasis | -0.786 | 0.100 | -1.166 | 0.120 |
|  | ERBB2 611CTF Targets | -0.959 | 0.200 | -0.757 | 0.280 |
|  | VEGF-A/VEGFR2 (WP) | -0.973 | 0.150 | -0.899 | 0.230 |
|  | HGF Signaling (AKT1-indep) | -1.307 | 1.000 | -1.337 | 1.000 |
|  | mTOR Signaling Down | -1.336 | 1.000 | -0.675 | 1.000 |
| Osimertinib | EGFR Signaling 24h Up | 1.761 | 0.251 | 1.535 | 0.091 |
|  | KRAS Lung Cancer Up | 1.987 | 0.125 | 2.066 | 0.005 |
|  | MYC Regulated Down | 2.084 | 0.054 | 2.536 | 0.002 |
| | NF-κB Fibroblast Targets | 1.686 | 0.286 | 2.224 | 0.003 |
|  | Apoptosis (Reactome) | 1.943 | 0.143 | 1.443 | 0.579 |
|  | Hypoxia Up | 1.986 | 0.119 | 2.215 | 0.003 |
|  | EGF Interval Up | -0.819 | 0.100 | -0.731 | 0.090 |
|  | Cell Cycle S Phase | -0.650 | 0.130 | -0.981 | 0.140 |
|  | Cell Cycle G1 Peaks | -0.651 | 0.100 | -1.211 | 0.170 |
| | IFN-$\alpha/\beta$ Signaling | -0.885 | 0.050 | -2.104 | 0.046 |
|  | MAPK Signaling (KEGG) | 1.011 | 0.089 | 2.169 | 0.003 |
|  | PI3K-AKT (WP) | 1.240 | 0.030 | 2.538 | 0.002 |
| Crizotinib | NF-κB Keratinocyte Targets | 2.006 | 0.020 | 2.326 | 0.002 |
|  | NRF2 Up | 2.137 | 0.018 | 2.394 | 0.001 |
|  | MYC Bound Genes | 2.452 | 0.003 | 2.244 | 0.001 |
|  | P53 Network (WP) | 2.299 | 0.007 | 1.704 | 0.039 |
|  | Cell Cycle (Reactome) | 2.104 | 0.016 | 2.217 | 0.001 |
|  | MET Signaling (Reactome) | -0.749 | 0.100 | -0.798 | 0.100 |
|  | MET Motility (Reactome) | -0.772 | 0.130 | -1.098 | 0.160 |
|  | EGF/EGFR Signaling (WP) | -0.780 | 0.090 | -0.731 | 0.130 |
|  | STAT3 Targets Down | -1.475 | 0.300 | -1.644 | 0.484 |
|  | ERBB Signaling (WP) | 0.984 | 0.605 | 1.497 | 0.105 |
|  | PI3K-AKT (Reactome) | 1.372 | 0.256 | 1.327 | 0.201 |
|  | PI3K-AKT (WP) | 1.529 | 0.151 | 1.752 | 0.029 |
| Adagrasib | KRAS Lung Cancer Up | -1.707 | 0.083 | -1.567 | 0.066 |
|  | Hypoxia Up | -2.374 | 0.005 | -2.028 | 0.028 |
|  | Doxorubicin Apoptosis Down | -2.011 | 0.030 | -1.964 | 0.036 |
|  | EMT-like Breast Cancer | -2.224 | 0.015 | -1.855 | 0.063 |
|  | MYC Oncogenic Targets | -1.851 | 0.051 | -2.166 | 0.026 |
|  | MAPK Signaling (KEGG) | -1.177 | 0.357 | -0.958 | 0.398 |
|  | MAPK Signaling (WP) | -1.317 | 0.257 | -1.010 | 0.555 |
|  | RAS Signaling (WP) | -1.621 | 0.116 | -0.996 | 0.165 |
|  | EGFR TKI Resistance (WP) | -1.616 | 0.117 | -1.556 | 0.172 |
|  | PI3K-AKT Cancer Signaling | -1.503 | 0.160 | -1.174 | 0.311 |
|  | TCF-dependent WNT | 0.789 | 0.160 | 0.764 | 0.180 |
|  | ERBB2/CDC25A Tumorigenesis | 0.707 | 0.160 | 1.283 | 0.200 |

**Table S11.** Performance on the regime of zero-shot generalization to unprofiled drugs, as we progressively increase the injected knowledge sources. Knowledge variants are defined as: no knowledge, randomly initialized knowledge encoders; +MoleculeSTM, initialization from a pretrained MoleculeSTM checkpoint; +PrimeKG, further pre-training with PrimeKG entities; +MAP-KG, additional pre-training with MAP-KG entities.

| Variant | HVG |  |  | Top-50 DEG |  |
| --- | --- | --- | --- | --- | --- |
| | Pearson $\Delta$ | PDS | Direction Acc | Pearson $\Delta$ | Direction Acc |
| no KG | $0.196 \pm 0.009$ | $0.055 \pm 0.003$ | $0.556 \pm 0.012$ | $0.321 \pm 0.011$ | $0.553 \pm 0.015$ |
| +MolSTM | $0.484 \pm 0.011$ | $0.136 \pm 0.005$ | $0.694 \pm 0.016$ | $0.698 \pm 0.015$ | $0.831 \pm 0.019$ |
| +PrimeKG | $0.591 \pm 0.013$ | $0.301 \pm 0.007$ | $0.742 \pm 0.018$ | $0.708 \pm 0.017$ | $0.862 \pm 0.022$ |
| +MAP-KG | **$0.659 \pm 0.014$** | **$0.344 \pm 0.006$** | **$0.807 \pm 0.021$** | **$0.822 \pm 0.023$** | **$0.934 \pm 0.024$** |

**Table S12.** Ablation of knowledge types by removing one type of knowledge entities drom MAP-KG at a time, including drug-side textual attributes (*text*: name/MOA/therapeutic description), gene-side attributes (*seq*: amino-acid sequence; *text*: functional annotations), and relational edges (drug–gene mechanistic interactions and gene–gene functional/physical associations), under the regime of zero-shot generalization to unprofiled drugs.

| Model | HVG |  |  | Top-50 DEG |  |
| --- | --- | --- | --- | --- | --- |
| | Pearson $\Delta$ | PDS | Direction Acc | Pearson $\Delta$ | Direction Acc |
| w.o. gene text | $0.594 \pm 0.018$ | $0.271 \pm 0.026$ | $0.735 \pm 0.018$ | $0.690 \pm 0.013$ | $0.870 \pm 0.010$ |
| w.o. gene-gene edges | $0.610 \pm 0.027$ | $0.292 \pm 0.019$ | $0.740 \pm 0.020$ | $0.709 \pm 0.023$ | $0.881 \pm 0.024$ |
| w.o. drug text | $0.567 \pm 0.025$ | $0.216 \pm 0.023$ | $0.733 \pm 0.002$ | $0.704 \pm 0.010$ | $0.868 \pm 0.011$ |
| w.o. drug-gene edges | $0.558 \pm 0.009$ | $0.231 \pm 0.017$ | $0.719 \pm 0.014$ | $0.679 \pm 0.012$ | $0.854 \pm 0.012$ |
| MAP-KG | **$0.659 \pm 0.014$** | **$0.344 \pm 0.006$** | **$0.807 \pm 0.021$** | **$0.822 \pm 0.023$** | **$0.934 \pm 0.024$** |

### A.2 Detailed functional analysis on A-549

In Section 2.4, we performed pathway-level evaluation by applying GSEA to MAP’s zero-shot predictions on held-out drugs in A549, and showed that the resulting ranking prioritizes four clinically established NSCLC therapies. Here we complement this ranking with drug-by-drug functional interpretations by inspecting representative gene sets (pathways) that are linked to each compound’s known mechanism of action and to disease-relevant programs in A549.

#### Afatinib

Afatinib is an irreversible pan-ERBB inhibitor [45]. In Figure 4c, MAP closely matches the experimental GSEA on inflammatory and stress-associated programs, including NF-*κ*B signaling–related pathways, with highly concordant NES values. This agreement is biologically plausible given the prominent role of NF-*κ*B–mediated survival and inflammatory transcription in lung adenocarcinoma contexts [46, 47], and indicates that the model captures treatment-associated remodeling of these core programs. In addition, MAP reproduces a modest activation of apoptosis-related pathways, consistent with drug-induced cellular stress responses observed in the measured perturbation profiles.

#### Crizotinib

Crizotinib is a clinically used kinase inhibitor in NSCLC [48]. Figure 4c shows that MAP accurately recovers the enrichment of NRF2-associated oxidative stress response pathways, consistent with the well-characterized KEAP1–NRF2 antioxidant-defense axis in A549 [49]. The model also matches the enrichment of cell-cycle–related programs, indicating that it preserves proliferative regulatory states reflected in the transcriptomic data. Together, the concurrent agreement on NRF2 and cell-cycle gene sets supports that MAP yields context-consistent and mechanistically interpretable pathway readouts for this compound.

#### Adagrasib

Adagrasib is a covalent KRAS^G12C^-directed therapy designed to attenuate downstream RAS– MAPK signaling [50, 51]. In the A549 perturbation data, key KRAS-associated oncogenic programs exhibit clear negative enrichment, and MAP closely reproduces these suppressive patterns. Notably, the model correctly predicts downregulation of KRAS-driven lung cancer signatures, accompanied by reduced activity of downstream proliferative programs such as MYC-associated gene sets. This coordinated agreement forms a coherent pathway-level response that is consistent with attenuation of KRAS-linked transcriptional output observed in the experimental GSEA.

#### Osimertinib

Osimertinib is a third-generation EGFR inhibitor [52]. In Figure 4c, MAP reproduces the experimentally observed activation of an EGFR signaling response signature, consistent with acute pathway perturbation and compensatory/feedback regulation captured in the transcriptomic profiles. At the same time, the model matches persistent enrichment of KRAS-related oncogenic programs and NF-*κ*B–associated inflammatory pathways, indicating that it correctly preserves dominant disease-relevant transcriptional programs in this cellular context [53, 54]. Overall, these results demonstrate that MAP can recover drug-proximal responses while maintaining biologically coherent background regulation at the gene-set level.

### A.3 Evaluation Metrics

**Pearson delta correlation** calculates Pearson correlation coefficient(PCC) between predicted and observed log fold changes(logFC) in gene expression, and quantifies how well the model recovers the magnitude of perturbation-induced expression changes. Formally, given a perturbed pseudo-bulk *x*^(*p*)^, the model prediction 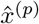 and a paired control pseudo-bulk *x*, the Pearson Delta Correlation is calculated as

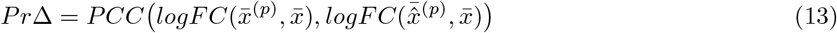

where 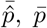 and 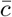 denote the cell-wise mean expression of corresponding pseudo-bulks, and PCC(,) is calculated as:

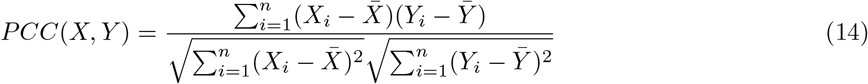

The coefficient ranges from -1 to 1, where 1 indicates perfect positive correlation, -1 indicates perfect negative correlation, and 0 indicates no linear relationship. We calculate *Pr*Δ for both highly variable genes (HVG) and top-50 differentially expressed (DE) genes.

**Direction accuracy** evaluates whether the model predicts the correct direction of regulation relative to the control state, and is formulated as:

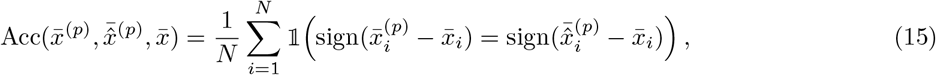

where *N* is the number of evaluated genes (HVGs or top-50 DE genes), *x*_*i*_ ∈ ℝ denotes the expression value of *i*-th gene, *sign*(·) denotes the sign of the input value and 𝟙 (·) denotes the indicator function

**Perturbation discrimination score** assesses whether the predicted perturbation effects are distinguishable across different perturbations. Given *T* perturbations, we define for each perturbation *p*:

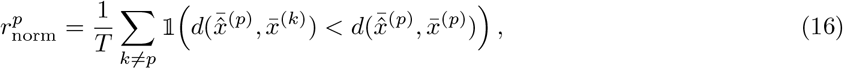

where *d*(·, ·) is the Manhattan distance. The perturbation discrimination score is defined as:

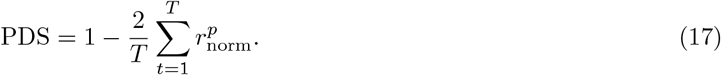

We calculate perturbation discrimination score only on HVGs to ensure consistency across perturbations.

### A.4 Details of Knowledge Sources

To ensure the diversity, high quality and broad coverage of MAP-KG, we curate and integrate fragmented knowledge from 14 public sources. Here, we briefly introduce each source below, and summarize their contributions of knowledge data in Table S13.

**BindingDB [55]** is a public database specializing in experimentally measured binding affinities of small molecules to their protein targets.

**BioGRID [56]** (Biological General Repository for Interaction Datasets) is a curated biological database of protein, chemical, and genetic interactions. It archives interaction data from multiple organisms, supporting network-based analysis of cellular pathways and gene function.

**ChEMBL [57]** is a manually curated, open-source database of bioactive molecules with drug-like properties, maintained by the European Bioinformatics Institute (EMBL-EBI). It provides a vast repository of high-quality bioactivity data extracted from scientific literature and deposited datasets.

**DrugBank [58]** is a bioinformatics and cheminformatics resource that combines detailed drug data with comprehensive drug target information. It integrates chemical, pharmacological, and pharmaceutical data with detailed target sequence and pathway information, making it a gold-standard database for pharmacologists and pharmaceutical researchers.

**DrugCentral [59]** is a comprehensive online drug database that provides information about active pharmaceutical ingredients. It serves as a key resource for drug discovery and development by integrating drug data from various authoritative sources, including information on indications and pharmacological actions.

**Ensembl [36]** is a comprehensive genome database that provides integrated access to high-quality annotated genome sequences for vertebrates and other model organisms. It aggregates genomic and gene annotation data, including gene models, transcript and protein sequences, variation and comparative genomics information, and provides tools for data retrieval and analysis to support a range of genomic research tasks.

**HGNC [37]** (HUGO Gene Nomenclature Committee) database is the authoritative resource for standardized human gene nomenclature. It assigns unique symbols and names to each human gene to ensure clear scientific communication and facilitate unambiguous data retrieval from publications and other databases.

**Human PPI [60]** is a computational resource that provides a predicted human protein-protein interactome. It allows users to search for interactions via gene names or UniProt identifiers and to check for interactions between specific protein pairs.

**IntAct [61]** is an open-source database system that provides a freely available resource for molecular interaction data. It offers a rich platform for studying protein functions in cellular processes through experimentally determined interactions.

**PrimeKG [34]** is a biomedical knowledge graph that integrates curated information from public resources into a unified graph. It links entities such as diseases, drugs, genes/proteins, pathways/biological processes, phenotypes, and anatomical concepts. We incorporate the drug–gene relationships from this resource into MAP-KG.

**PubChem [35]** is a public repository of chemical molecules and their biological activities, maintained by the National Center for Biotechnology Information (NCBI). It provides extensive data on chemical structures, identifiers, physicochemical properties, and bioassay results, serving as a foundational resource for chemical biology and drug discovery research.

**STITCH [62]** (Search Tool for Interacting Chemicals) is a database of known and predicted interactions between chemicals and proteins. It integrates interaction data from experiments, databases, and literature to build comprehensive networks.

**STRING [63]** is a biological database and web resource for known and predicted protein-protein interactions. These interactions include both direct (physical) and indirect (functional) associations.

**TTD [64]** (Therapeutic Target Database) is a specialized resource that provides information about known therapeutic protein and nucleic acid targets, along with their targeted diseases, pathway information and the corresponding drugs directed at each of these targets.

**Table S13.**
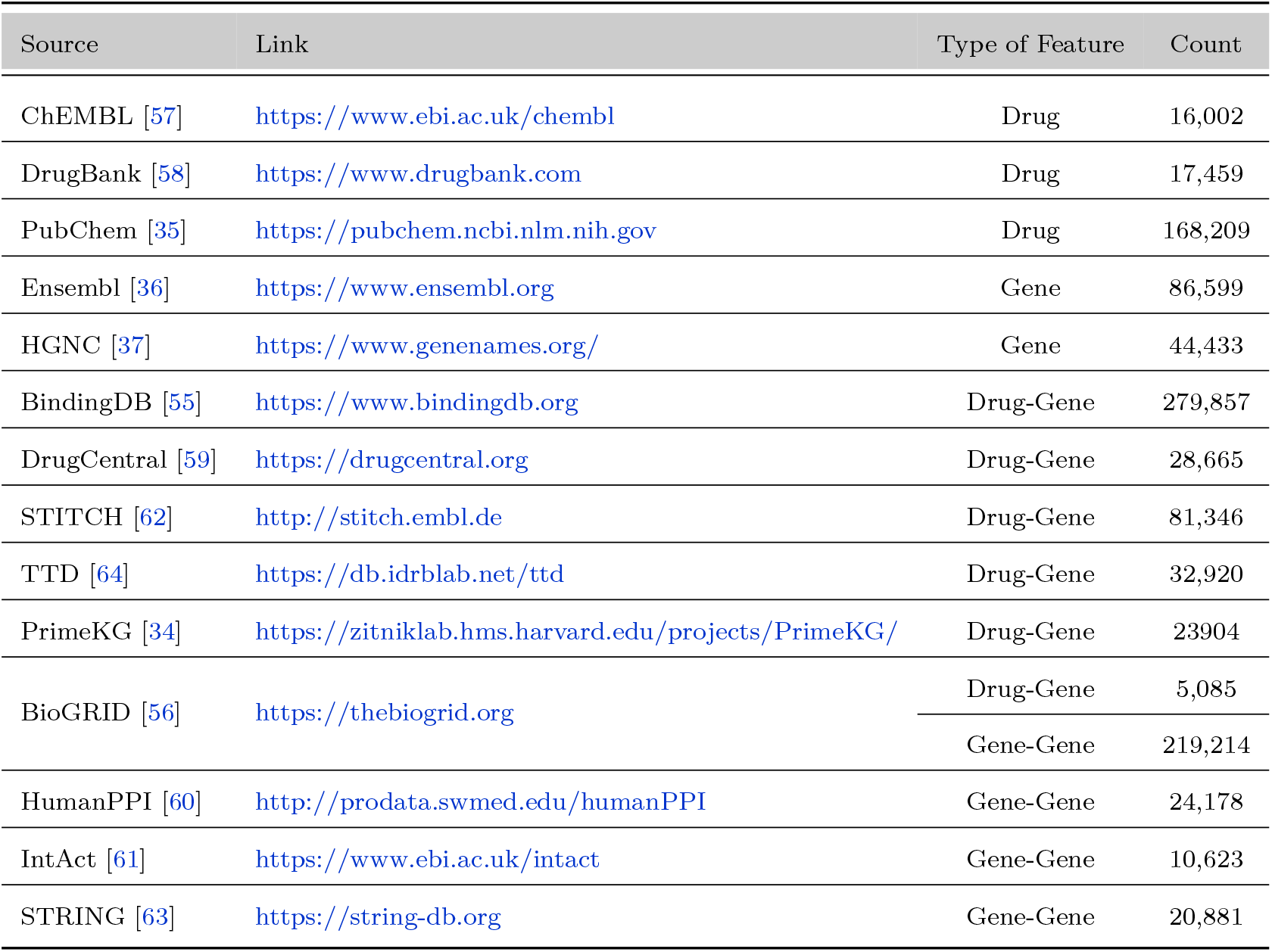
Links and statistics of the 14 knowledge sources in MAP-KG.

**Table S14.**
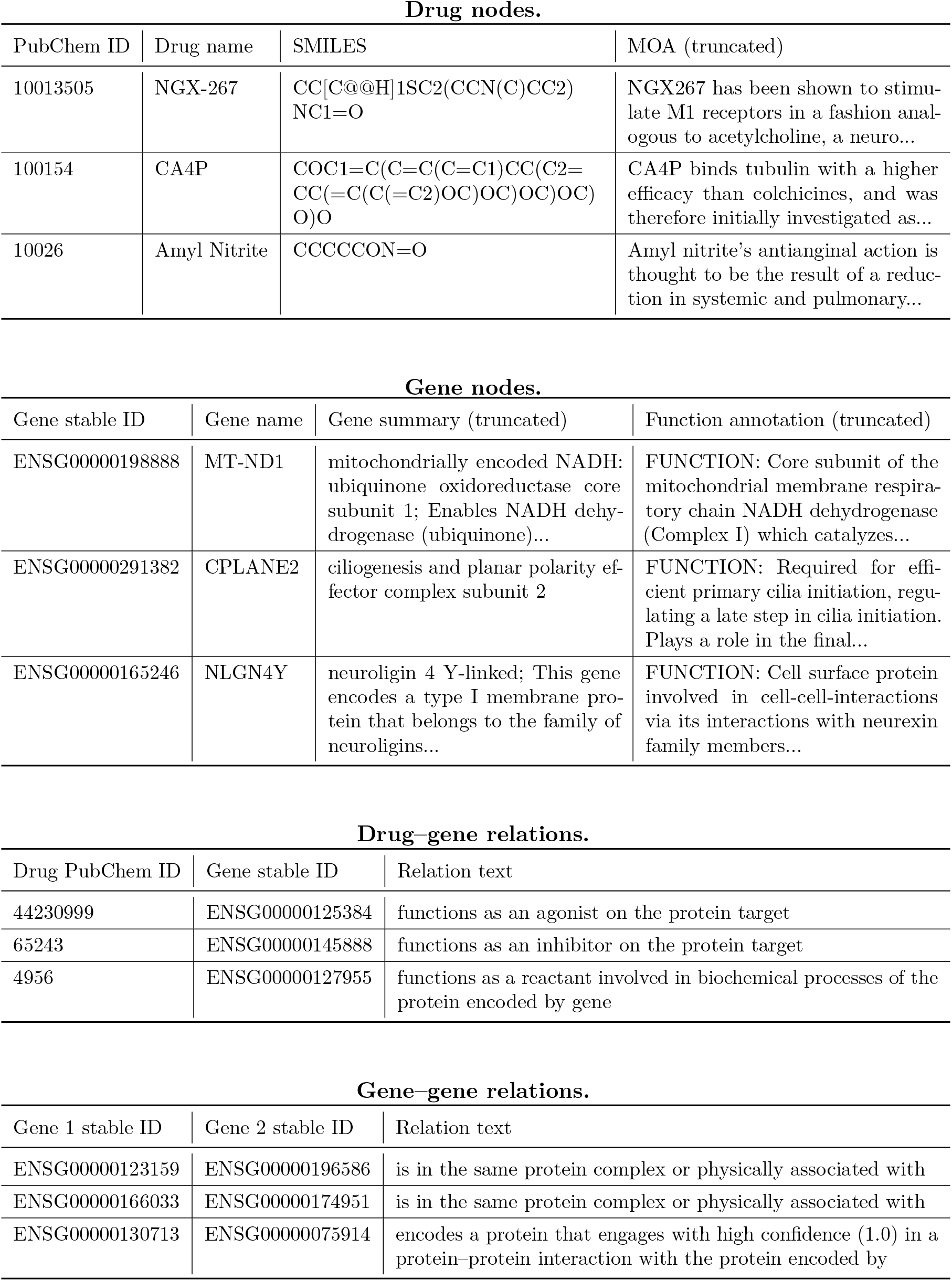
Representative examples of nodes and edges in MAP-KG. We list example drug nodes, gene nodes, and relation edges with the attributes used in our graph construction. Long text fields are truncated to the first 25–40 words for readability. Canonical gene sequences are included as attributes in MAP-KG but omitted here due to length.

